# Probing the interaction between the WDR41 7CD loop of the C9ORF72 complex and purified PQLC2

**DOI:** 10.64898/2026.09.18.752389

**Authors:** L Jamal, P von Gaudecker, L Pieri, C Ransy, N Nakamae, F Andre, I Tascon, H Jiang, B Ochoa-Lizarralde, I Ubarretxena-Belandia, C Debacker, B Gasnier, LC Tabares, JL Vázquez-Ibar

## Abstract

The lysosomal cationic amino acid transceptor PQLC2 integrates transport with receptor-like signaling roles by recruiting the C9ORF72/SMCR8/WDR41 (CSW) complex in response to cellular cationic amino acid depletion. CSW recruitment depends on the interaction of PQLC2 with a disordered loop (7CD loop) extending from the WDR41 β-propeller. Defining the molecular basis of this interaction is therefore essential for understanding how PQLC2 connects cellular amino acid availability to downstream signaling. Here, we used site-directed electron paramagnetic resonance (EPR) spectroscopy to characterize the interaction between purified PQLC2 and a synthetic WDR41 7CD loop peptide containing a site-specific nitroxide spin label. Using a consensus mutagenesis strategy, we stabilized detergent-purified PQLC2 and found that it assembles as a homotrimer, similar to other PQ-loop transporters. EPR experiments demonstrated the direct and specific association of the 7CD loop with PQLC2, and highlighted the key role of the the central region of the 7CD loop to establish this interaction via conserved aromatic residues. Spin labels introduced at different positions further provided local structural and dynamic information of the 7CD loop upon PQLC2 binding. The data supports a model in which the central region of the loop is anchored within the cytosolic cavity of PQLC2, whereas the flanking regions, which are dispensable for binding, retain substantial conformational flexibility. Together, we provide a direct biochemical evidence for PQLC2/WDR41 7CD loop recognition, a step forward towards understanding the transceptor mechanism of PQLC2.

## Introduction

Lysosomes are membrane-bound organelles responsible for degrading and recycling cellular waste and macromolecules (1,2). In addition to their role in cellular catabolism, the lysosomal membrane also functions as a dynamic signaling platform for nutrient sensing and metabolic signal transduction (2–5), where membrane-embedded lysosomal membrane transport proteins play essential roles, particularly on nutrient sensing (6). The best characterized lysosomal-mediated cell signaling pathway involves the multiple protein complex mTORC1 (mechanistic target of rapamycin complex 1), a critical controller in cellular growth, survival, metabolism, and development (7). mTORC1 integrates signals such as cellular nutrient availability (e.g., amino acids) and triggers responses that stimulate anabolism or suppress catabolism (3,8–10). Recruitment and activation of mTORC1 occurs at the lysosomal surface and is mediated by the Rag GTPases whose nucleotide loading status is regulated by amino acid availability (11–13). Biochemical and structural data have uncovered the role of the lysosomal amino acid transceptor SNAT9/SLC38A9 on sensing amino acid content leading to mTORC1 activation (14–18). SNAT9 mediates the efflux of essential amino acids from lysosomes, such as leucine, in an arginine-regulated manner (14,19). In the absence of substrate, the N-terminus of SNAT9 forms a β-hairpin that inserts into the cytoplasmic open cavity of the transporter, near the substrate-binding pocket (18,20). When arginine is abundant, binding of arginine to the transporter releases the N-terminal plug (20), which then interacts with the Rag-GTPase complex switching their nucleotide state to activate mTORC1 (17,18). Plug release also indirectly activates mTORC1 by enabling SLC38A9 to export other essential amino acids, which stimulate other cytosolic amino acid sensors (17,18,20).

Recent studies have revealed an amino acid sensing role for another lysosomal amino acid transporter, the PQ-loop repeat containing protein 2 (PQLC2/SLC66A1), involved in a less characterized lysosomal-mediated cell signaling pathway (21,22). PQLC2 is responsible of exporting cationic amino acids from the lysosomal lumen to the cytosol (23,24). In addition to its transporter function, PQLC2 recruits a protein complex composed by C9ORF72, SMCR8, and WDR41 (also known as CSW complex) to the lysosomal surface under cellular cationic amino acid starvation (21,22,25). PQLC2 is thus a ‘transceptor’ (transporter-like receptor) (26) since it combines both, a transporter and a receptor function. The CSW complex is involved in several lysosome-mediated functions, including autophagy, mTORC1 or Toll-like receptors regulation (27–31). Importantly, a hexanucleotide repeat expansion in a non-coding region of the C9orf72 gene is a major cause of two neurodegenerative diseases: amyotrophic lateral sclerosis (ALS) and frontotemporal dementia (FTD) (32,33). While the atomic 3D structure of PQLC2 is not yet determined, Cryo-EM studies have revealed the molecular architecture of the CSW complex (22,34). Moreover, “*in vitro*” biochemical studies have shown that the CSW complex display a C9ORF72, SMCR8-mediated GTPase-activating protein (GAP) activity to a specific set of small GTPases, that includes Rab8A, Rab11A, Arf1, Arf5 and Arf6 (22,34). Interestingly, none of these GTPases are localized at the lysosome, suggesting that CSW lysosomal recruitment during cationic amino acid starvation may serve, among other potential roles, as a regulatory mechanism for its GAP activity. PQLC2 consists of 7 predicted transmembrane-spanning domains (TM), with the C- and N-termini located, respectively, in the cytoplasm and in the lysosomal lumen (24). High-resolution structures of several members of the PQ-loop family have brought to light the 3D fold architecture of this family, consisting in two symmetrically-related triple helix bundles (TMs 1–3 and TMs 5-7) connected by the TM 4 (35–38). Homodimers of bacterial semi-SWEET transporters share also the same structural 3D fold (39,40). A large solvent-accessible cavity where the substrate binding site is located opens to either side of the membrane, allowing substate translocation across the membrane (37,41).

Pull-down and co-localization studies performed in cellular cultures have determined that the recruitment of the CSW complex by PQLC2 occurs exclusively through the interaction between PQLC2 and WDR41 (21,22,25). Importantly, this interaction is reversible since replenishment of cationic amino acids in the cell culture medium induces the release of the CSW complex from the lysosomal membrane (21,22,25). WDR41 consists of an eight bladed β-propeller structure and interacts with SMCR8 through its C-terminal helix domain (22,34). Using mutagenesis and pull-down assays, Ferguson and colleagues identified a critical loop in WDR41—connecting blade 7 between β-strands C and D (7CD loop)—as the determinant domain that mediates the interaction with PQLC2 (25). Specifically, a 10-amino-acid motif within the “turn” sequence of the 7CD loop (‘Transporter-Interacting Peptide’ or TIP sequence) seems to be sufficient to mediate interaction with PQLC2 during lysosomal recruitment of the CSW complex (25). The 7CD loop cannot be modeled in any of the existing cryo-EM structures of the CSW complex (22,34). This finding suggests an inherent flexibility and a lack of defined secondary structure of this loop, also observed by hydrogen/deuterium exchange mass spectrometry studies (22).

The interaction between PQLC2 and the 7CD loop of WDR41 triggered by cellular cationic amino acid depletion is critical for the recruitment of the CSW complex to the lysosomal membrane, and its subsequent signaling activity. So far, the mechanism by which PQLC2 senses the low amino acid content to bind the 7CD loop is still unknown. To gain deeper insights into the transceptor role of PQLC2, in this work we sought to define the molecular basis of the interaction between PQLC2 and the WDR41 7CD loop using purified protein components and site-directed electron paramagnetic resonance (EPR) spectroscopy. We first applied a consensus mutagenesis approach to engineer stability of purified PQLC2 in detergent micelles. We demonstrated the direct association between the 7CD loop peptide and purified PQLC2, and probed the critical role of the TIP motif via conserved aromatic residues. We then mapped the local structural dynamics of the 7CD loop by introducing nitroxide spin labels at different positions along the peptide. These analyses provide residue-specific structural information of the 7CD loop upon binding to PQLC2, supporting a model in which the central TIP motif is tightly anchored within the cytosolic cavity of PQLC2, whereas the flanking regions, which are dispensable for binding, remain conformationally flexible.

## Results

### Expression and purification of a mammalian PQLC2

We utilized the yeast *Saccharomyces cerevisiae* as expression host to produce recombinantly the rat ortholog of PQLC2 (rPQLC2) (which shares 86% amino acid identity with human PQLC2 (Figure 1)) fused to the superfolder green fluorescent protein (GFP) at the C-terminal end, and including an engineered Hrv-3C protease recognition site between the GFP and the C-terminus of rPQLC2. Previous studies have demonstrated that rPQLC2 is functional in *S. cerevisiae* as it complements the canavanine sensitivity phenotype of YPQ2, a yeast homolog of PQLC2 (24,42). YPQ2 activity enhances canavanine toxicity by exporting the drug from the vacuolar lumen to the cytoplasm. Expression of rPQLC2 in a *ypq2* knockout mutant restores the YPQ2-dependent canavanine sensitivity phenotype, confirming that yeast-expressed rPQLC2 is capable of transporting canavanine (24,42).

**Figure 1.**
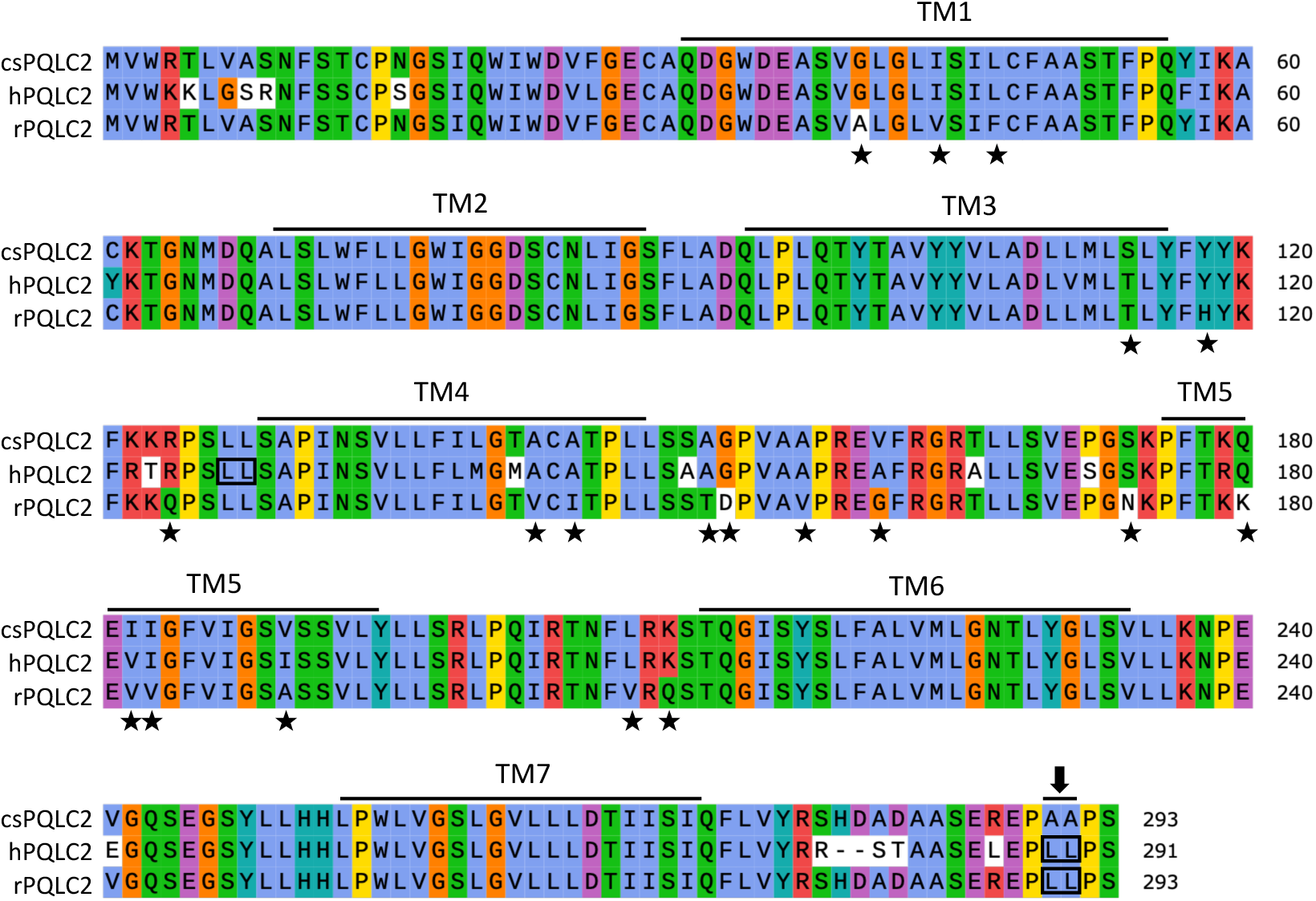
Multiple sequence alignment of human, rat, and consensus-based engineered PQLC2 sequences. csPQLC2: consensus-based engineered rPQLC2; hPQLC2: human PQLC2 (Uniprot accession number Q6ZP29); rPQLC2: rat PQLC2 (Uniprot accession number B0BMY1). Amino acid identity between the hPQLC2 and rPQLC2 is 86%. Amino acid identity between the rPQLC2 and csPQLC2 is 93%. Residues replaced in rPQLC2 to build the csPQLC2 sequence are indicated by (★). Lines above define the predicted transmembrane (TM) domains of rPQLC2 calculated using TOPCONS (Tsirigos et al., 2015) (Bernsel et al., 2009). The dileucine sorting motifs are framed. In the csPQLC2 sequence, the C-terminal dileucine motif were replaced by alanine (indicated by an arrow). Residues are colored according the Clustal X Colour Scheme The alignment was performed by Muscle.

A common strategy to redirect lysosomal membrane transporters to the plasma membrane consists in mutating their lysosomal sorting motif, often located at the N- or C-terminal end of the transporter (24,43,44). In *Xenopus* oocytes, mutating the di-leucine based sorting motif in rPQLC2 by alanine (L290A/L291A, for simplicity referred as LL/AA) results in the redirection of the GFP-tagged rPQLC2 from the lysosomal membrane to the plasma membrane (24). Similar mutations of two di-leucine based sorting motifs in human PQLC2 (Figure 1) expressed in HEK293 cells also redirects the transporter to the plasma membrane (25). With this in mind, we expressed in *S. cerevisiae* two versions (wild type and LL/AA) of the GFP-tagged rPQLC2.

Consistent with previous reports (24,42,45), fluorescence microscopy confirmed that rPQLC2-GFP localizes almost exclusively in the membrane of the vacuole, the analogue of the mammalian lysosome (Figure 2A). Interestingly, mutation of the di-leucine sorting motif to alanine (LL/AA) did not alter the membrane localization of rPQLC2-GFP (Figure 2A), in contrast to previous observations (45). Western-blot analysis of isolated membranes using an anti-GFP antibody revealed a single band around 50 kDa, consistent with the theoretical molecular weight of rPQLC2-GFP (∼59 kDa) (Figure 2B).

**Figure 2.**
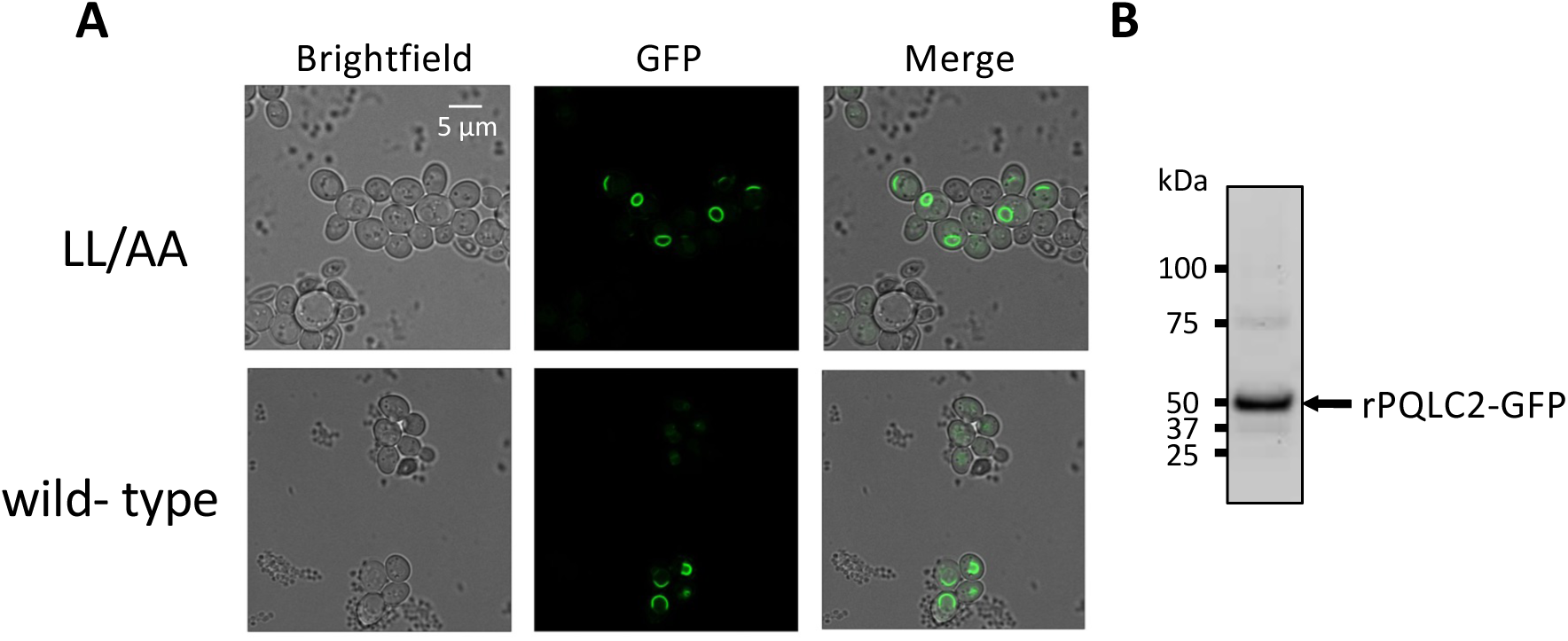
Expression of rPQLC2-GFP in *S. cerevisiae*. (*A*) Fluorescent confocal microscopy images of *S. cerevisiae* cells expressing either LL/AA or wild type rPQLC2-GFP. The images show that both versions of PQLC2-GFP are located in enclosed vacuole organelles. (*B*) Western blot of membranes expressing LL/AA rPQLC2-GFP using the antibody against the GFP.

Since the LL/AA version of rPQLC2 showed a better expression yield than the wild type, we decided to continue with this version for the subsequent solubilization and purification tests. In addition, LL/AA rPQLC2 has already been shown to be fully functional in *Xenopus laevis* oocytes (24,46). Membranes expressing LL/AA rPQLC2-GFP were solubilized with dodecyl maltoside (DDM) supplemented with cholesteryl hemisuccinate (CHS), and protein monodispersity in detergent micelles was analyzed by fluorescence size-exclusion chromatography (FSEC) (47) (Supplementary Figure S1A). The FSEC chromatogram of DDM-solubilized LL/AA rPQLC2-GFP showed a monodisperse profile, confirming the suitability of DDM/CHS to efficiently solubilize LL/AA rPQLC2-GFP from the membranes. LL/AA rPQLC2 was purified by incubating the detergent-solubilized membranes with agarose beads coupled to an anti-GFP nanobody (GFP-Trap) (48), followed by cleavage with the HRV-3C protease. A single band at ∼25 kDa in SDS-PAGE confirmed the purity of the sample (Figure 3A). Mass spectrometry analysis further confirmed that this electrophoretic band corresponds to the expected rPQLC2 sequence (Figure 3B). Although our purification protocol yielded a pure protein sample in reasonable quantities (∼1.5 mg of purified PQLC2 per liter of yeast culture), size-exclusion chromatography (SEC) revealed that LL/AA rPQLC2 exhibited poor stability in detergent micelles at concentrations ≤ 1 mg mL^-1^. We also purified wild-type rPQLC2 using the same strategy. However, wild-type rPQLC2 showed even less stability in solution than the LL/AA version as it aggregated during concentration, preventing further SEC analysis.

**Figure 3.**
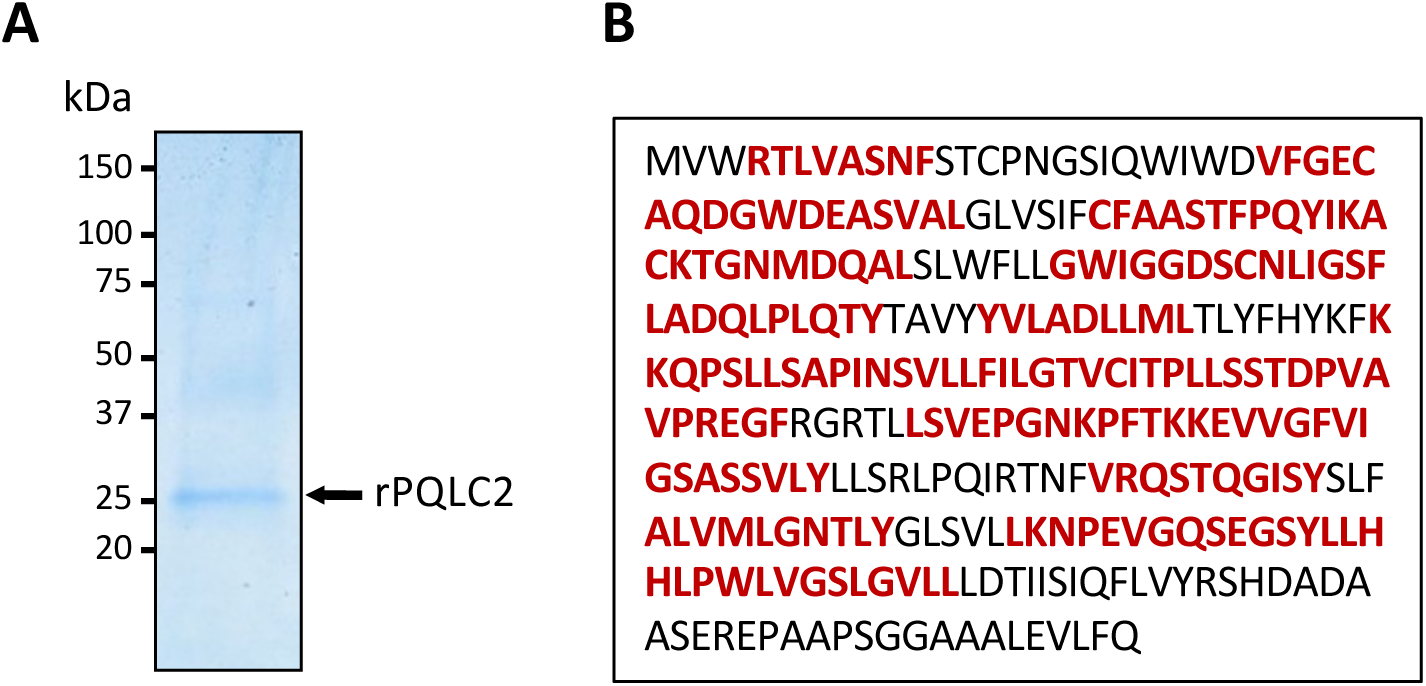
Characterization and identification of purified LL/AA rPQLC2. (**A**) Coomassie blue staining SDS-PAGE of affinity-purified LL/AA rPQLC2 expressed in *S. cerevisiae* (**B**) the rPQLC2 electrophoretic band of (*A*) was analyzed by LC-MS/MS after chymotrypsin digestion. The peptides identified by mass spectrometry are displayed in red, and cover 68% of the entire sequence.

### Engineering stability in purified rPQLC2

As obtaining pure and stable PQLC2 is critical to study “*in vitro*” its interaction with the 7CD loop, we employed a protein engineering approach based on consensus mutagenesis to enhance the stability of purified LL/AA rPQLC2 in detergent micelles. This method has been shown to effectively improve the thermal stability of membrane proteins post-purification (49,50). We adopted a strategy similar to that used by Reyes and collaborators to thermostabilize the human excitatory amino acid transporter EAAT1 (49). We first generated a consensus amino acid sequence based on the alignment of 22 homologous PQLC2 sequences (Supplementary Figure S2). Residues in the rPQLC2 sequence with low conservation were replaced by the corresponding residue of the consensus sequence, provided that the consensus residue occurred with a frequency ≥ 20% and was at least 10% more frequent than the original residue in rPQLC2. The resulting engineered LL/AA rPQLC2 sequence (referred to as csPQLC2) shares 93% amino acid identity with rPQLC2, and contains 19 substitutions, primarily located in the predicted transmembrane regions and the loop connecting TMs 4 and 5 (Figure 1). Despite the relatively low sequence conservation of both the N- and C-terminal ends among PQLC2 homologs (Supplementary Figure S2), we decided to not introduce any substitutions in these regions. This is because motifs involved in membrane trafficking—such as the di-leucine motif—are often present in these terminal regions (43,44).

To confirm that the amino acid substitutions introduced in csPQLC2 did not compromise transporter functionality, we expressed the LL/AA versions of rPQLC2-GFP and csPQLC2-GFP in *Xenopus laevis* oocytes. Epifluorescence microscopy revealed that rPQLC2-GFP and csPQLC2-GFP exhibit comparable expression levels at the plasma membrane of the oocytes (Figure 4A-B). We then measured rPQLC2-mediated [³H] L-arginine uptake, an established assay for characterizing the functional properties of rPQLC2-GFP (24,46). In these assays, csPQLC2-GFP transported [³H] L-arginine into oocytes to a similar extent as the non-engineered version (Figure 4C), indicating that the introduced amino acid substitutions did not compromise the transport acitvity. Additionally, fluorescence microscopy confirmed that csPQLC2-GFP also localizes to the vacuole membrane when expressed in *S. cerevisiae* (Supplementary Figure S3). We purified the yeast-expressed csPQLC2-GFP in DDM/CHS using the same protocol as for rPQLC2-GFP. Unlike the non-engineered version, size-exclusion chromatography (SEC) analysis of the protein concentrated to ∼50 µM (1.7 mg mL^-1^) revealed a homogeneous elution profile with minimal aggregation (Supplementary Figure S4A). This confirms that csPQLC2 exhibits enhanced stability in DDM/CHS detergent micelles compared to the original sequence, thus enabling further biophysical analysis in detergent micelles.

**Figure 4.**
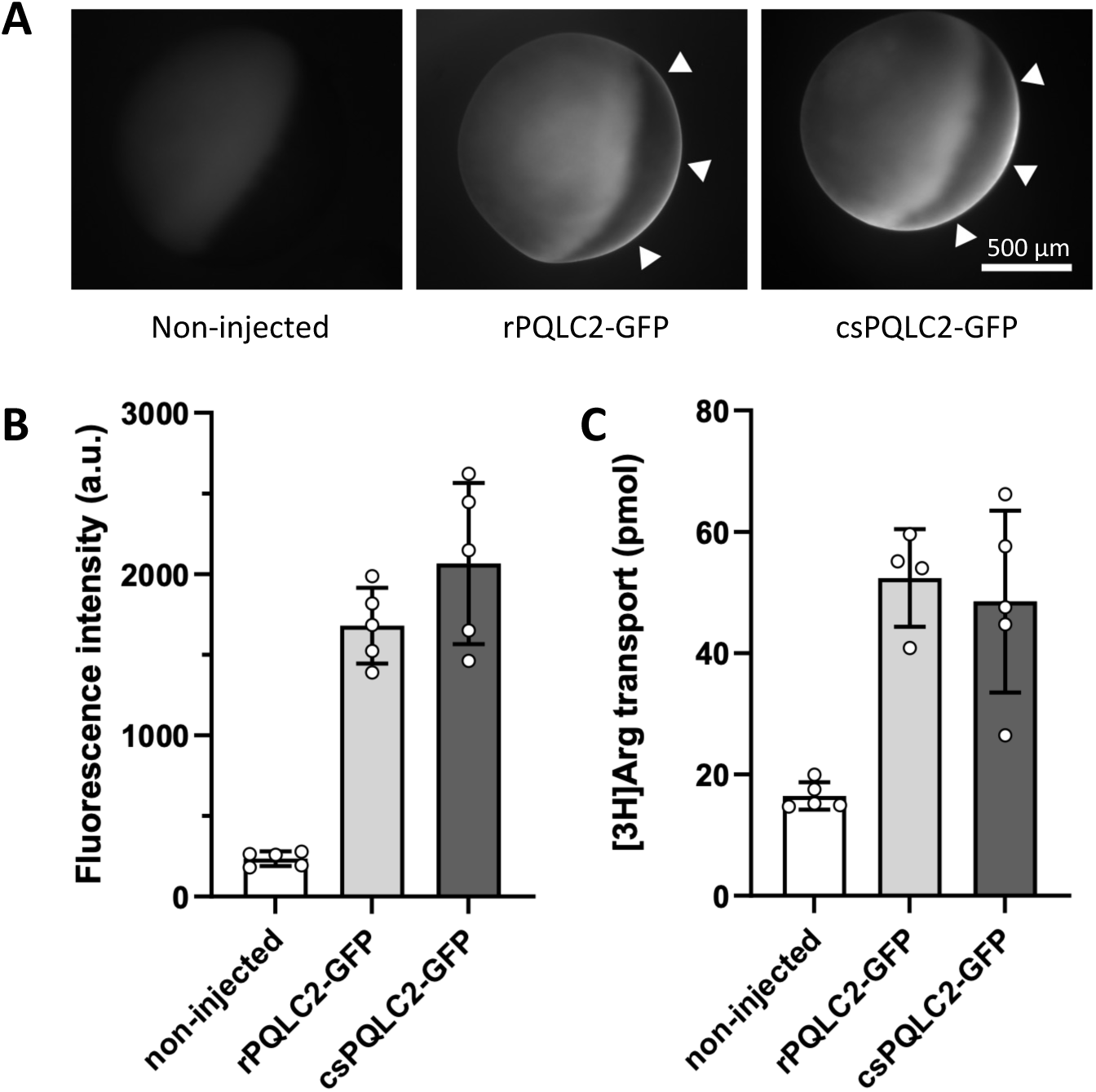
Expression and functional assessment of PQLC2 constructs in *Xenopus* oocytes. (A) The expression of rPQLC2-EGFP or csPQLC2-EGFP at the plasma membrane of *Xenopus oocytes* (indicated by arrows) was analyzed by epifluorescence microscopy using the GFP fluorescence as reporter. (B) GFP fluorescence was quantified at the plasma membrane. Each value represents the mean ± SEM of five oocytes per condition. (C) Uptake of [^3^H] L-Arg into oocytes expressing rPQLC2-GFP or csPQLC2-GFP. Transport was initiated by adding [^3^H]L-Arg (100 *μ*M; 0.5 *μ*Ci) at pH 5.0 and stopped after 20 min by brief ice-cold washes. Accumulated radioactivity was counted individually for each oocyte. The mean ± SD of five oocytes per condition is shown.

### Purified csPQLC2 adopts a homo-oligomeric organization

Some members of the PQ-loop family are known to assemble as homotrimers (35,51). The retention time seen in both FSEC and SEC chromatograms suggests also potential oligomerization of PQLC2 in detergent micelles (Supplementary Figures S1 and S4), although SEC analysis is not accurate enough to assess the oligomerization state of detergent-solubilized membrane proteins. Using AlphaFold v2.0, we generated a structural model of trimeric rPQLC2 (Figure 5A). All three protomers in this trimeric model are almost identical (Root Mean Square Deviation (RMSD) ∼ 0.04 Å). Excluding the N- and C-terminal regions and the loop connecting TMs 4 and 5, the predicted local distance difference test (pLDDT) or confidence scores for this model range between 60 and 90 for most residues of the trimer, indicating moderate to high confidence (Supplementary Figure S5). In this model, each rPQLC2 protomer adopts the characteristic architecture of PQ-loop transporters, consisting of two triple-helix bundles (TMs 1–3 and TMs 5–7) connected by TM4 (Figure 5A-B) (35,37,51). Contacts between monomers are provided between TMs 3 and 4 of one protomer and TMs 5 and 7 of another protomer (Figure 5A). Importantly, the ipTM score of the rPQLC2 trimeric model, which measures the accuracy of the relative positions of the PQLC2 monomers, was 0.83 (scores above 0.8 indicate high-confidence predictions)(52). Overall, these scores support a reasonable level of confidence in the trimeric arrangement of PQLC2 predicted by AlphaFold.

**Figure 5.**
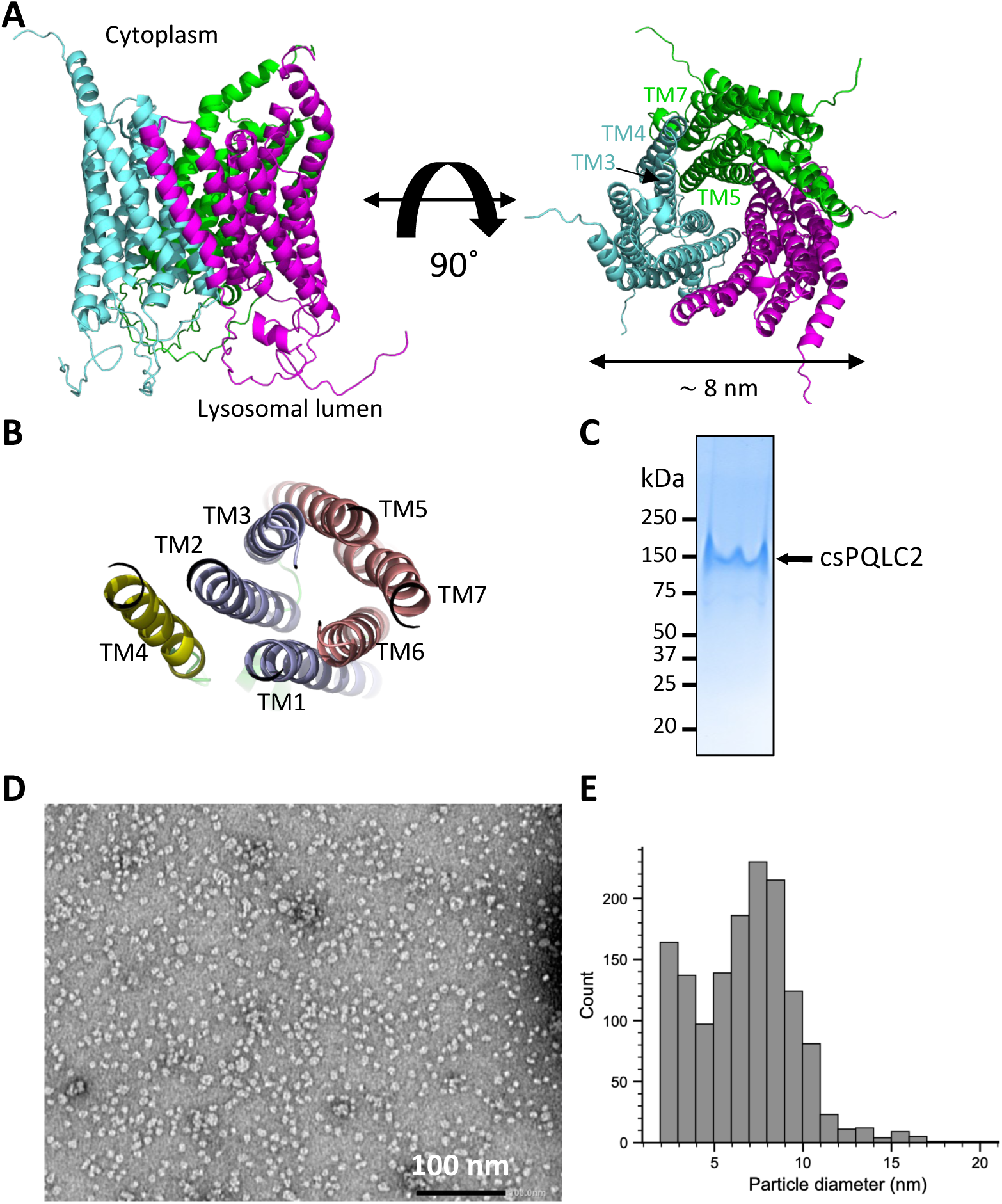
Oligomerization of PQLC2. (*A*) Cartoon representation of the rPQLC2 AlphaFold V2.0 trimeric model in lateral (left), and in the cytoplasmic-faced (right) orientation. Each monomer is represented by a different color. The TMs involved in protomer-protomer contact are indicated. (*B*) Cytoplasmic-faced conformation of the AlphaFold V2.0 model of monomeric rPQLC2. TMs 1, 2 and 3 (light blue) and TMs 5, 6 7 (salmon) form two similar helix bundles connected by TM4. Loops connecting TMs were removed from the picture. (*C*) Non-denaturing native gel electrophoresis of purified csPQLC2. (*D*) Representative NS-EM micrograph of purified csPQLC2 in 0.015 % (v/v) LMNG. (*E*) Histogram representation of particle diameter distribution of (*D*).

To investigate the oligomeric state of PQLC2, we conducted negative-staining electron microscopy (NS-EM) studies of detergent-purified csPQLC2. Unfortunately, NS-EM images of csPQLC2 in DDM/CHS were indistinguishable from the detergent background, consistent with previous observations (53). Consequently, we opted to exchange detergents from DDM to LMNG, as FSEC analysis had confirmed the efficacy of LMNG/CHS in solubilizing rPQLC2-GFP (Supplementary Figure S1B). Initial attempts to exchange detergents (from 0.02% (v/v) DDM to 0.015% (v/v) LMNG) on the GFP-Trap beads were unsuccessful, as no protein was eluted following Hrv-3C protease incubation. We then performed detergent exchange during SEC. csPQLC2 was solubilized and purified in DDM/CHS, concentrated, and injected into an SEC column equilibrated with 0.015% (w/v) LMNG. In these conditions, csPQLC2 eluted as a single, homogeneous peak (Supplementary Figure S4B). NS-EM analysis of the freshly SEC eluted csPQLC2 revealed particles of relatively uniform size (Figure 5D). The larger population of particles displayed diameters in the range of 8–10 nm (Figure 5E), consistent with the dimensions of the rPQLC2 trimeric model predicted by AlphaFold (∼8 nm) (Figure 5A). In addition, we found a subpopulation of particules in the range of 2-3 nm (Figure 5E) consistent with the predicted monomeric dimensions of rPQLC2. Purified csPQLC2 displayed an electrophoretic mobility closer to ∼100 kDa in a non-denaturing PAGE (Figure 5C), also in agreement with a homotrimeric arrangement of csPQLC2 (theoretical Mw of csPQLC2 homotrimer: 99330 Da). Together, AlphaFold prediction, NS-EM and biochemical data are consistent with a homotrimeric organization of detergent-purfied csPQLC2.

### WDR41-7CD loop interacts with purified csPQLC2

The interaction between the WDR41 7CD loop and PQLC2 has been proposed to be essential and sufficient to recruit the CSW complex to the lysosomal membrane (25). This interaction is suggested to occur via insertion of the TIP motif of the 7CD loop into the central cavity of the cytosol-open conformation of PQLC2 (25). This model is also supported by AlphaFold prediction (Figure 6C), that situates as well the TIP region of the 7CD loop at the bottom of the central cavity of the cytosolic-open conformer of PQLC2. To validate this hypothesis and elucidate the molecular basis of 7CD loop/PQLC2 recognition, we used room temperature EPR spectroscopy to investigate the interaction between purified csPQLC2 and a synthetic peptide corresponding to the 7CD loop (Figure 6). The proline at position 8, which plays no role in the interaction with PQLC2 (25), was mutated to cysteine and used as an anchoring point for a nitroxide radical via a maleimide reaction, generating the 7CD-P8x peptide (Figure 6). This spin label serves as a reporter of peptide dynamics. The EPR spectrum of a nitroxide radical in solution is strongly dependent on its rotational dynamics. Rapid molecular rotation produces three sharp lines, whereas restricted motion leads to progressive line broadening (54). This motion is typically described by the rotational correlation time (*τ*_c_), which can be obtained by spectral simulation. For a spin label, the calculated *τ*_c_ reflects local dynamics, global tumbling, and the degree of motion of the bonds connecting the nitroxide moiety to the peptide or protein. The latter effect is generally dominant. Therefore, *τ*_c_ provides a quantitative measure that represents the overall nitroxide mobility in the ∼100 ps to 10 ns time range (55).

**Figure 6.**
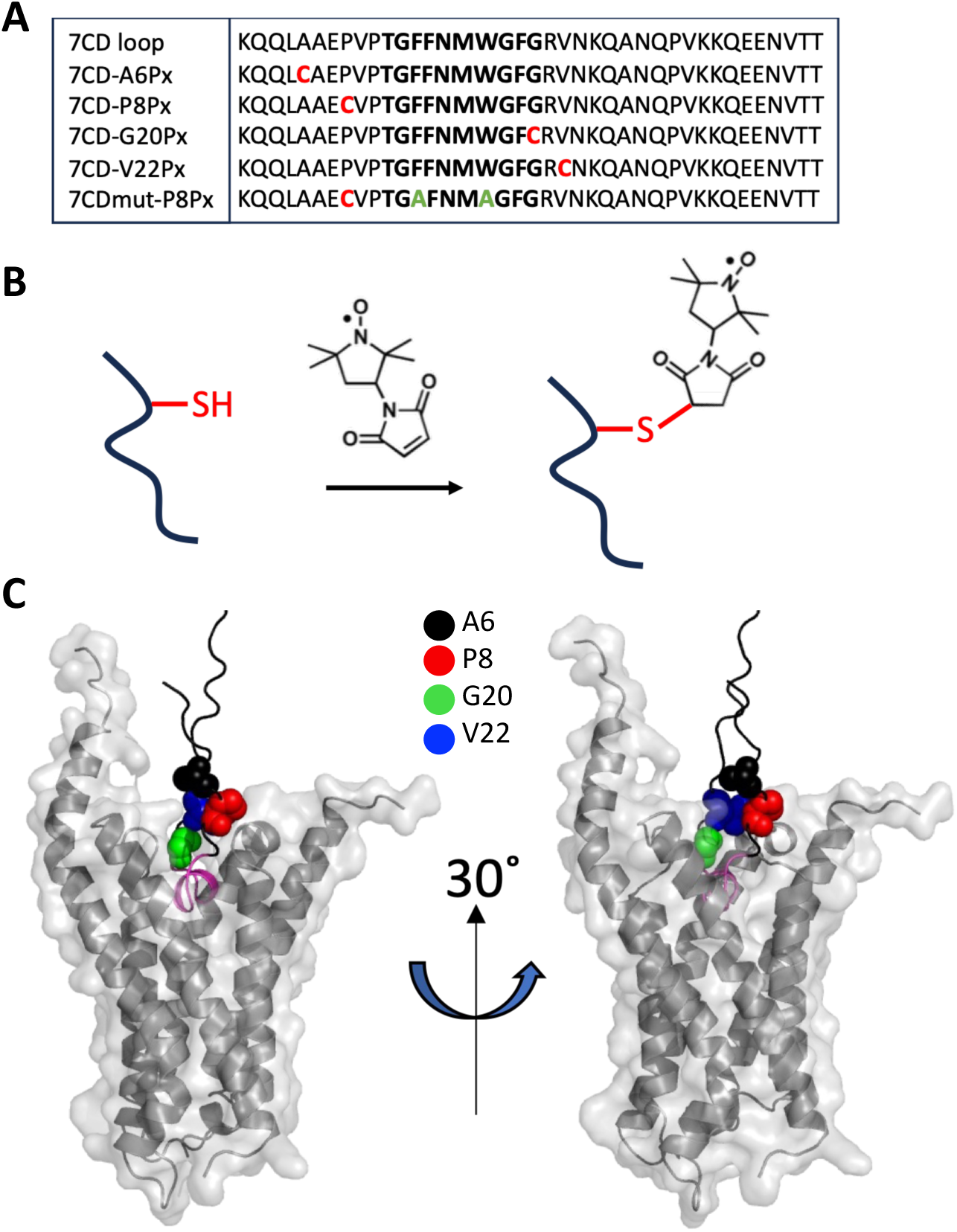
Spin-labeled 7CD-loop peptides variants and Alphafold V2.0 WDR41-7CD loop/csPQLC2 model. (*A*) Table showing the sequence of the different peptides used for the EPR studies. Bold letters indicate the TIP region. The red color highlights the position of the Cys residue introduced for labeling, and the green color indicates the residues replaced by Ala in the WDR41mut peptide. (*B*) Schematic representation of the reaction between the 3-maleimido-PROXYL spin label and the cysteine introduced at various positions of the 7CD peptide, resulting in the labeled peptides indicated in the top table. (*C*) AlphaFold V2.0 model of rPQLC2 (cartoon and surface representation) in complex with WDR41-7CD loop (cartoon). The WDR41-7CD loop TIP region is depicted in pink. The sphere representation in WDR41-7CD loop correspond to the positions where the nitroxide group was introduced, as indicated in the figure. For clarity, the TM4 of PQLC2 was removed from the model.

The line shape of the 7CD-P8Px EPR spectrum (Figure 7, top, black trace) is indicative of a fast-moving nitroxide, with a *τ*_c_ of ∼0.6 ns (see below), as expected for a small peptide. Upon addition of an equimolar amount of purified csPQLC2, a new feature appeared, clearly visible in the low-field region of the spectrum (Figure 7, top, red trace). Such a signal is characteristic of strongly restricted nitroxide motion (*τ*_c_ > 10 ns). We therefore attribute this slow-mobility component to a bound state in which the spin label directly interacts with csPQLC2, resulting in a drastic reduction in its mobility freedom. Further evidence was obtained from EPR measurements in frozen solution at 50 K. Low-temperature EPR allows determination of the hyperfine interaction along the z direction (*A_z_*) by direct measurement from the spectra or via simulations (Figure 7C and Table 1). *A_z_* correlates with the dielectric constant surronding the nitroxide spin label, varying from ∼95 MHz in organic solvents to ∼105 MHz in aqueous solutions (56). For 7CD-P8Px in the working buffer containing DDM, the *A_z_* value was 105 MHz, decreasing to 95 MHz in the presence of csPQLC2. This value is consistent with the label being located in a protein environment (25). To determine whether this interaction is driven by the specific interaction of the peptide with the protein, a competition experiment was performed by adding a five-fold molar excess of unlabeled 7CD loop. This resulted in a significant decrease in the slow-mobility component, indicating that 7CD-P8Px and the unlabeled 7CD loop compete for the same binding site (Figure 7A-B, green trace). The residual slow component at a 5:1 ratio of unlabeled to labeled peptide suggests that the label itself does not significantly affect the binding affinity. To confirm that this interaction is driven by the TIP motif, residues F13 and W17 were mutated to alanine in 7CD-P8Px (7CDmut-P8Px, Figure 6). Previous studies have shown that mutation of these residues (corresponding to residues F366 and W370 on full-length WDR41) abolishes PQLC2-mediated recruitment of the CSW complex (25). Consistently, the EPR spectrum of 7CDmut-P8Px was unaffected by the presence of csPQLC2 (Figure 7A-B), indicating that the mutated peptide does not interact with the transporter.

**Figure 7.**
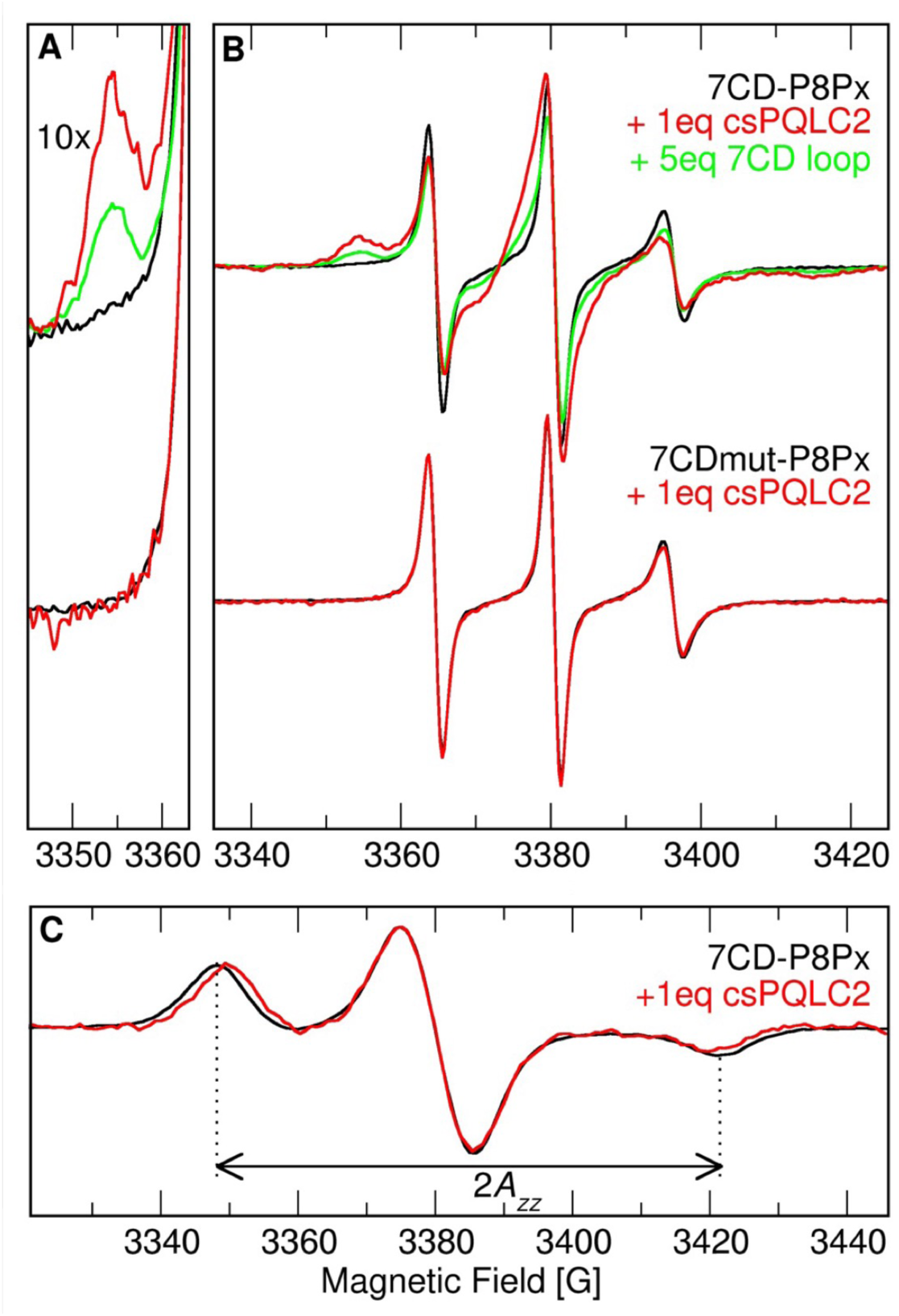
Room-temperature, 9 GHz EPR spectroscopy reveals the direct interaction between the 7CD loop and purified csPQLC2. (*B*) The spectra of 7CD-P8Px in buffer alone (black trace), after addition of 1 eq of csPQLC2 (red trace), and after subsequent addition of 5 eq of unlabeled 7CD loop (green trace). (Bottom) The spectra of 7CDmut-P8Px in buffer alone (black trace), after addition of 1 eq of csPQLC2 (red trace). Panel *A* shows a 10x expansion of the low-field, 3345-3363 G, region of the spectra. (*C*) 50K 9 GHz spectra of 7CD-P8Px in buffer alone (black trace), after addition of 1 eq of csPQLC2 (red trace). All spectra have been normalized to the unity of their double integral.

**Table 1-.** EPR fitting parameters. The different spectra of the spin-labeled peptides shown in Figure 8, were fitted in the absence (no csPQLC2) or presence (+ 1eq. csPQLC2) of purified csPQLC2. A single component model was used in the experiments in the absence of csPQLC2 and in the ones with the 7CDmut-P8Px peptide, whereas a three-components model was used in the rest of the experiments.

| Component | Parameter | 7CD-A6Px | 7CD-P8Px | 7CD-G20Px | 7CD-V22Px | 7CDmut-P8Px |
| --- | --- | --- | --- | --- | --- | --- |
| **No csPQLC2** |  |  |  |  |  |  |
| **Unbound** | **$t_c$ (ns)** | **0.52** | **0.64** | **0.88** | **0.83** | **0.55** |
|  | Axy (MHz) | 15.0 | 15.0 | 14.9 | 14.7 | 15.0 |
|  | Az (MHz) | 104.1 | 105.0 | 104.3 | 104.0 | 104.6 |
|  | rmsd | 0.009 | 0.006 | 0.011 | 0.018 | 0.010 |
| **+ 1eq. csPQLC2** |  |  |  |  |  |  |
| **Unbound** | **$t_c$ (ns)** | **0.51** | **0.62** | **0.80** | **0.85** | **0.64** |
|  | Axy (MHz) | 13.3 | 13.4 | 13.2 | 13.2 | 15.0 |
|  | Az (MHz) | 107.5 | 106.9 | 106.9 | 107.5 | 104.0 |
|  | Weight | 0.36 | 0.24 | 0.19 | 0.31 | 1.000 |
| **Bound-Restricted** | **$t_c$ (ns)** | **11.34** | **13.42** | **22.25** | **22.09** |  |
|  | Axy (MHz) | 16.8 | 14.5 | 14.2 | 16.0 |  |
|  | Az (MHz) | 95.4 | 96.0 | 96.4 | 95.4 |  |
|  | Weight | 0.33 | 0.40 | 0.30 | 0.23 |  |
| **Bound-Dynamic** | **$t_c$ (ns)** | **2.34** | **3.50** | **4.81** | **2.97** |  |
|  | Axy (MHz) | 13.7 | 14.4 | 13.1 | 15.3 |  |
|  | Az (MHz) | 99.8 | 100.2 | 103.1 | 103.6 |  |
|  | Weight | 0.30 | 0.36 | 0.50 | 0.46 |  |
|  | rmsd | 0.004 | 0.003 | 0.008 | 0.003 | 0.007 |

To gain further structural and dynamic insights into the 7CD loop/PQLC2 interaction, room-temperature EPR experiments were repeated with the spin label localized at upstream (A6 and P8), and downstream (G20 and V22) positions of the TIP region (Figure 6A-C). All peptides behaved similarly to 7CD-P8Px, showing a slow-mobility component only in the presence of csPQLC2 (Figure 8A-B). However, its relative intensity strongly depended on the label position, being highest for 7CD-G20Px, located near the C-terminal end of the TIP region, and decreasing with increasing distance from the TIP, reaching a minimum for 7CD-A6Px (Figures 6 and 8).

**Figure 8.**
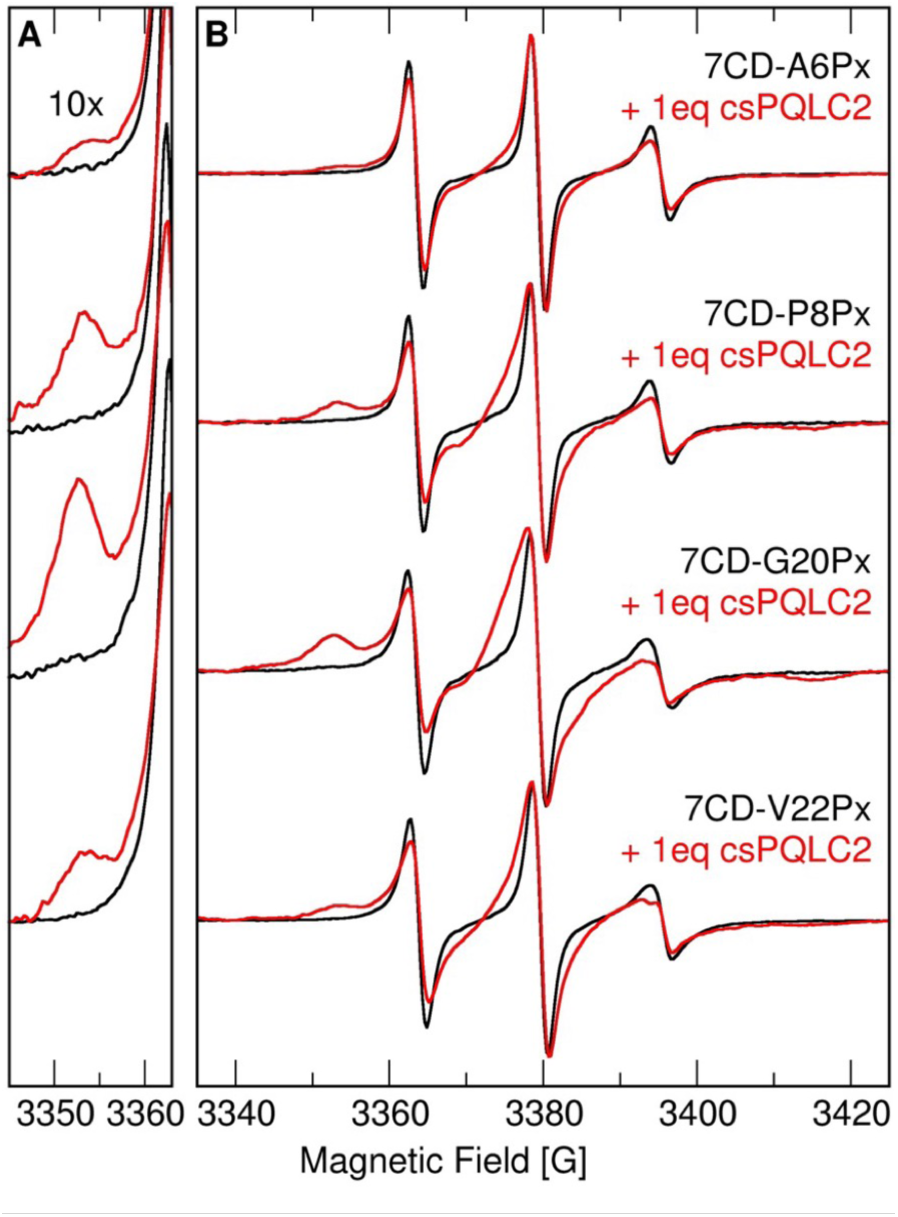
Room-temperature, 9 GHz spectra of the 7CD loop labeled at different positions. (*B*) Spectra of 7CD labeled at the positions indicated in the figure in buffer alone (black traces), and after addition of 1 eq of csPQLC2 (red traces). A 10x expansion of the low-field, 3345-3363 G, region of the spectra is shown in panel *A*.

To quantify these observations, the spectra were simulated. In the absence of csPQLC2, all peptides were well described by a single component with rotational *τ*_c_ in the 0.5–0.8 ns range, depending on the labeling position, likely reflecting subtle structural differences (Table 1, Supplementary Figure S6). Consistent with its inability to interact with csPQLC2 (Figure 7), the spectrum of 7CDmut-P8Px in the presence of csPQLC2 was also accurately simulated using a single component with rotational *τ*_c_ of 0.5 ns (Figure 7A-B, Table 1). For the other peptides in the presence of csPQLC2, an initial simulation using a two-component model was performed. However, the resulting fits were not satisfactory, and systematic deviations were observed in the residuals (Table 1, Supplementary Figure S7A). Consequently, a three-component model comprising a fast, an intermediate, and a slow component was then employed. In this model, the initial values for fitting *τ*_c_ were set either to the values obtained for the free peptides (fast component) or to match the low-field resonance feature (slow component), and were then allowed to vary within a small range. The *τ*_c_ of the intermediate component was set to an intermediate value and allowed to vary over a larger range (see Supporting Information for details). This approach significantly improved the quality of the fits and substantially reduced the residual error (Table 1, Supplementary Figure S7B), therefore indicating that at least three populations coexist in equilibrium (Table 1). The fast component corresponds to the unbound peptide (unbound, Table 1), whereas the slow component likely arises from peptides bound to csPQLC2 in a conformation where the spin label directly interacts with the protein, resulting in the strongest restricted mobility (bound-restricted, Table 1). The intermediate component reflects a population of peptides bound to csPQLC2, but in a state where the spin label retains substantial motional freedom, and therefore not restrained to a single conformation on the timescale of the EPR experiment (bound-dynamic, Table 1). It should be noted, however, that this component may also arise from a fast exchange between multiple conformations, resulting in an apparent intermediate mobility. The coexistence of bound-dynamic and bound-restricted populations could reflect peptide binding to distinct conformational or oligomeric states of csPQLC2 in solution, including the trimeric and monomeric forms (Figure 5). However, a second scenario where the TIP region remains anchored to a unique binding site, while the rest of the 7CD peptide chain explores different conformations in the same binding site, is also possible. This second scenario is indeed appealing as 7CD-G20Px, containing the spin label near the C-term of the TIP region (Figure 6 A-C), display the largest *τ*_c_ of the bound components, thus reflecting the most restricted mobility among all labeled positions (Table 1). Conversely, 7CD-A6Px in which the label remains more exposed to the solvent away from the TIP region, presented the smallest *τ*_c_ values of the bound components (Table S1, Figure 6A-C). Molecular dynamics (MD) simulations of the AlphaFold 7CD loop/csPQLC2 complex (Supplementary Figure S8) further supported this second scenario. The complex was embedded in a micelle composed of 200 DDM molecules, and two independent 0.5 μs simulations were carried out. The results showed that, while the TIP region remained fixed in the putative binding pocket of csPQLC2, the rest of the peptide was highly flexible. Analysis of the backbone mobility at the labeling positions (Supplementary Figure S8) showed that the Cα atoms explored conformational spaces that could account for the coexistence of the bound-dynamic and bound-restricted components observed in the EPR spectra, each of which may itself comprise more than one conformation. Moreover, we observed that, whereas the Cα atom at position G20 explored a volume of approximately 0.6 nm^3^ during the simulation, the Cα atom at position A6 explored a much larger volume of approximately 3.2 nm^3^ (Suplemented Figure S8). This is consistent with 7CD-G20Px displaying the largest *τ*_c_ values for both, the bound-dynamic and bound-restricted components, whereas 7CD-A6Px displayed the smallest values (Table 1). On the other hand, the peptides labelled at P8 and V22 exhibited intermediate behavior, consistent with their structural environments in the model (Figure 6A-C and Supplementary Figure 8).

Overall, these results supports a 7CD loop–PQLC2 inteaction model where the TIP motif, containing the essential residues F13 and W17, is anchored into the central cavity of the cytosol-open conformation of PQLC2 (Figure 8C), while the rest of the peptide, non essential for the interaction, retains a certain degree of flexibility while bound within PQLC2’s cavity.

## Discussion

PQLC2 has recently emerged as a lysosomal amino acid transceptor that combines a cationic amino acid transport activity with the recruitment of the CSW signaling complex to the lysosomal membrane in response to cellular cationic amino acid starvation (21,25). Since CSW’s recruitment is reversible upon amino acid replenishment, PQLC2 operates as cationic amino acid sensor, thus modulating the lysosomal-dependent CSW’s signaling cascade according to the cellular amino acid content.

Pull-down assays in cell extracts and immunofluorescence microscopy have shown that the recruitment of the CSW complex is specifically mediated by the interaction between the protruding 7CD loop of WDR41 and PQLC2 (21,57), where a conserved 10-amino acid motif (TIP) within this protruding loop plays a critical role (25). However, the molecular determinants of PQLC2-WDR41 recognition and modulation in the presence of amino acids remain poorly characterized, largely because no experimental high-resolution structures of PQLC2 alone or in complex with WDR41 are currently available. AlphaFold modeling of the PQLC2–7CD loop complex predicts that the TIP motif inserts into the central cavity of the cytosolic-facing conformation of PQLC2 (Figure 6C), in agreement with previous predictions combining homology modeling and molecular docking (25). In this work, to gain mechanistic insight into the role of PQLC2 as both an amino acid transporter and a signaling receptor, we investigated the interaction between purified PQLC2 and the WDR41 7CD loop peptide using site-directed electron EPR spectroscopy.

To obtain a purified mammalian PQLC2, we decided to produce the rat homolog of PQLC2 (rPQLC2) in the yeast *S. cerevisiae* since it is expressed exclusively in the vacuole membrane (Figure 2), and previous studies demonstrated that rPQLC2 functionally complements the canavanine-sensitive growth phenotype of its yeast homolog, YPQ2, indicating that the recombinant protein is correctly folded and retains its transport activity (24,42). However, contrary to previous findings (Llinares et al., 2015), we observed that, in our experimental conditions, the conserved C-terminal dileucine-based sorting motif (Figure 1) is not mandatory for vacuole sorting in *S. cerevisiae* (Figure 2). This discrepancy might reflect the distinct culture conditions used in the two studies.

Purified rPQLC2 showed poor stability in detergent micelles, incompatible for downstream experiments. To overcome this, we employed a consensus-based mutagenesis strategy. This protein engineering approach has recently proven effective in enhancing the thermal stability of various membrane transporters, thereby facilitating their structural and functional analysis (49,50,58). The success of this approach is based on experimental observations showing that amino acids that appear more frequently at a given position in a sequence have a stronger stabilizing effect than those appearing less often (59). The engineered csPQLC2 demonstrated significantly improved stability compared to wild-type, as evidenced by its monodisperse behavior in size-exclusion chromatography (SEC) at concentrations suitable for EPR experiments (Supplementary Figure S4). Importantly, functional assays in *Xenopus* oocytes confirmed that csPQLC2 transports L-Arg similarly to the wild-type non-engineered version (Figure 4). Analysis of detergent-purified csPQLC2 by NS-EM and native PAGE indicated that csPQLC2 likely assembles as homotrimer, a finding supported as well by AlphaFold modeling (Figure 5 and Supplementary Figure S5). Interestingly, the crystal structure of another member of the PQ-loop family, the vacuolar glucose transporter OsSWEET2b (35) display a similar trimeric architecture as the one predicted by Alphafold for rPQLC2 (Figure 5B). Although the recent structures of human and plant cystinosin transporters, the closest homologs of PQLC2, appear monomeric (36,37), homooligomeric organization has been observed in other PQ-loop family members, such as the vitamin B3 transporter PnuC (51) or the *Arabidopsis* SWEET sugar transporters (60,61). Notably, oligomerization seems crucial for function in some plant SWEET transporters, suggesting potential allosteric coupling between protomers during substrate translocation (61). Future studies combining high-resolution structural data and functional assays will be necessary to determine whether PQLC2 homooligomerization influences its transport activity or CSW receptor function.

Site-directed spin labeling combined with EPR spectroscopy is a powerful tool for probing molecular interactions and dynamics of membrane proteins in solution (55,62). In this work, we have exploited this approach to investigate the association between purified csPQLC2 and the 7CD loop peptide using a nitroxide spin label as reporter, strategically attached to non-essential positions of the 7CD loop (Figures 6, 7 and 8). The detection of a specific low-mobility EPR component of the spin-labeled peptide at room temperature, as well as low-temperature measurements, confirmed a direct association between the purified transporter and the 7CD loop (Figure 7). Importantly, EPR also proved the essential role of two conserved aromatic residues within the TIP region to establish this interaction (Figures 6 and 7A-B), corroborating previous pull-down studies (25). Mutation of these residues abolishes PQLC2 binding without disrupting WDR41 association with SMCR8 (25); therefore, our results are consistent with these findings, indicating that the TIP region specifically mediates PQLC2 recognition and plays no role on CSW complex integrity.

EPR analysis of spin labels introduced at different positions of the 7CD loop flanking the TIP region (Figures 6 and 8) provided position-specific insights into the local environment and conformational dynamics of the 7CD loop upon binding to csPQLC2 (Figure 8A-B). These experiment reveled that positions closer to the TIP motif exhibited the highest restriction in mobility, whereas more distal residues retained more mobility (Figure 8A-B, Table 1). Analysis of both restricted and dynamic bound states in the EPR spectra (Table 1), together with MD simulations (Supplementary Figure S8), indicates that the TIP region is stably anchored within the PQLC2 binding-site, serving, as previously resported, as the primary determinant of binding (25). The remaining regions of the 7CD loop display higher conformational freedom, consistent with its limited contribution to binding specificity (25). Together, these findings support a model in which a rigid TIP-mediated anchoring of the 7CD loop is accompanied by dynamic positioning of the flanking peptide segments within the PQLC2 cavity.

In summary, this work provides new insights towards understanding the transceptor mechanism of PQLC2 by characterizing the molecular basis of PQLC2-WDR41 interaction using purified proteins and peptides. By combining EPR spectroscopy and in silico studies together with protein engineering tools, we provide biochemical evidence of the direct interaction between the WDR41 7CD loop and PQLC2, and confirmed that the conserved aromatic residues within the TIP region are key determinants of PQLC2-7CD loop recognition. Moreover, our EPR studies support a structural model of PQLC2–7CD loop complex in which the TIP motif serves as the key anchoring element that inserts on PQLC2’s cytosolic cavity, whereas the regions flanking the TIP motif not essential for binding, remain conformationally flexible. Further structural and functional studies will be required to fully elucidate the transceptor mechanism of PQLC2.

## Material and Methods

### Construction of the expression vector pYeDP60 PQLC2

The cDNA encoding rPQLC2 (either wild-type or L290A/L291A) was amplified by PCR from the pOX(+)-rPQCL2-EGFP plasmid (24), and cloned into the *EcoRI*/*NotI* sites of the pYeDP60 expression vector (63). A synthetic cDNAs encoding csPQLC2 was obtained from GenScript and inserted between the *EcoRI*/*NotI* sites of the pYeDP60 vector.

### Introduction of superfolder GFP

To generate the GFP-tagged construct, the gene encoding the superfolder GFP (hereafter referred to as GFP) was introduced as previously described (48). A PCR fragment encoding the GFP, flanked at the 5ʹ end by a human rhinovirus 3C protease recognition sequence (Hrv-3C or 3C protease) and by a stop codon at the 3ʹ end, was inserted into the pYeDP60-rPQLC2 vector between the *NotI* and *XmaI* restriction sites, yielding the final construct pYeDP60-rPQLC2-3C-GFP.

### Yeast transformation

The different versions of PQLC2 were expressed in the *Saccharomyces cerevisiae* strain W303.1b Gal4.2 (α, leu2, his3, trp1: TRP1-GAL10-GAL4, ura3, ade2-1, canr, cir+) (64). Yeast transformations were performed using the lithium acetate method (65). Briefly, 5 mL of S6AU minimal media [0.1% (w/v) Bacto^TM^ Casamino Acids, 0.7% (w/v) Yeast Nitrogen Base (without amino acids and with ammonium sulphate)], 20 μg mL^-1^ adenine and uracil, supplemented with 2% (w/v) glucose], was inoculated with one yeast colony of non-transformed W303.1b/Gal4.2 strain, and incubated for 24 h at 28°C with shaking at 180 rpm. 2 mL of the above culture were harvested by centrifugation (5 min, 5,000 × *g*), and the pellet was mixed with ∼1 μg of plasmid DNA and 100 μg of salmon sperm DNA (denatured at 100°C for 5 min). The mixture was vortexed, and 500 μL of 40 % (w/v) PEG 4000, 100 mM lithium acetate, 10 mM Tris-HCl pH 7.5, 1 mM EDTA, and 35 mM DTT was added. The tube was vortexed again and left overnight at room temperature. The next day, the cell suspension was centrifuged (5 min, 2,000 x *g*), and the pellet was resuspended in a small volume of the S6A selective medium (S6AU minimal media without uracil). The resuspended pellet was plated onto a S6A-agar plate supplemented with 2% (w/v) glucose, and incubated at 28°C for 3-5 days until the transformed colonies appeared.

### Protein expression in *S. cerevisiae*

Cultures were initiated by inoculating 5 mL of S6A medium with freshly transformed yeast colonies, followed by incubation for 24 h at 28°C, shaking at 180 rpm. 20 mL (small-scale) or 500 mL (large-scale) of YPGE2X medium (2% (w/v) BactoTM Peptone, 2% (w/v), yeast extract, 1% (w/v) glucose and 2.7% (v/v) ethanol) were inoculated with this preculture to reach an initial OD_600_ of 0.05. Cells were cultured for 36 h at 28°C, shaking at 130 rpm, in order to consume the majority of glucose. Then, the culture flasks were cooled down to 18°C, and 2% (w/v) of galactose was then added to initiate protein induction. After 13 h, a second addition of 2% (w/v) galactose was made, and cells were cultured for another 5 hours. Cells were harvested by centrifugation (10 min, 4,000 × *g*, 4°C), and the cell pellet was washed with Milli-Q® H_2_O. Cell pellets were resuspended in 2 mL of TEKS buffer (50 mM Tris-HCl pH 7.5, 1 mM EDTA, 0.1 M KCl and 0.6 M sorbitol) per gram of cell pellet and after a 15 min incubation at 4°C, they were centrifuged again and the resulting cell pellet was flash-frozen in liquid nitrogen and stored at −70°C.

### Membrane preparation

Frozen cell pellets were resuspended in ice-cold TES buffer (50 mM Tris-HCl pH 7.5, 1 mM EDTA and 0.6 M sorbitol), 1 mL per gram of cell pellet, supplemented with SigmaFast^TM^ EDTA-free Protease Inhibitor Cocktail (PIC) (Sigma-Aldrich) and 1 mM phenylmethylsulfonyl fluoride (PMSF, Sigma-Aldrich). Cells were broken using glass beads (0.5 mm diameter) in a Planetary Mono Mill PULVERISETTE 6 (Fritsch). The broken crude extract was recovered, and the beads were washed with 1.5 mL of ice-cold TES buffer per gram of cell pellet. The pH of the crude extract was adjusted to 7.5, and cell debris, unbroken cells, and glass beads (designated as P1) were removed by centrifugation (20 min at 1,000 x *g*, 4 °C). The resulting supernatant (designated as S1) was subjected to a second centrifugation (20 min at 20,000 x g, 4°C) to obtain the P2 membranes. The supernatant (designated as S2) was subjected to ultracentrifugation (1 h, 125,000 x *g*, 4°C) to obtain the P3 membranes. Both, P2 and P3 membranes were resuspended in 0.2 mL HEPES-sucrose buffer (20 mM HEPES-Tris pH 7.4, 0.3 M sucrose, 0.15 M NaCl) per gram of cell pellet, flash-frozen in liquid nitrogen and stored at −70 °C.

### Fluorescent confocal microscopy

1 mL of a small-scale fresh yeast culture was centrifuged (5 min at 2,000 × *g*) and washed twice with 1 mL PBS pH 7.4. The cells were resuspended in 1 mL of PBS pH 7.4, and 2 μL were deposited on top of an agarose pad (1% (w/v) agarose gel in Milli-Q® H_2_O), casted on a glass slide. A coverslip was then applied and sealed with nail polish. Fluorescent images were captured using a Leica TCS SP8 multiphoton microscope, equipped with an HC PL APO 63x/1.40 OIL objective, at excitation and emission wavelengths of 488 and 580 nm, respectively. Image acquisition and processing were performed using LasX and Fiji software, respectively. Confocal microscopy was conducted at the Light Microscopy Facility of the I2BC (Gif-sur-Yvette, France).

### Protein detection by western blotting

Total protein concentration of membranes was measured by the BiCinchoninic acidAssay (BCA). Samples were subjected to SDS-PAGE and proteins were transferred to Polyvinylidene difluoride (PVDF) membranes (Immobilon®-P, Merck). GFP-tagged to PQLC2 was detected with a primary mouse antibody IgG1 K Anti-GFP (Roche) diluted to 1:1000 in 5% (w/v) milk in PBS-Tween buffer, followed by a incubation with a second Goat Anti-Mouse IgG-HRP conjugate antibody (Bio-Rad) diluted to 1:3000 in 5% (w/v) milk in PBS-Tween buffer. The ECL Western Blotting Detection kit (GE healthcare) was used for western blot revelation, and luminescence was detected by a CDD camera (Syngene).

### Detergent screening and fluorescence-detection size-exclusion chromatography (FSEC)

P3 membrane fractions were adjusted to a final protein concentration of 5 mg mL^-1^ in the solubilization buffer (20 mM Tris-HCl pH 7.8, 150 mM NaCl, 10% (v/v) glycerol, 5 mM MgCl₂, supplemented with PIC, and 1 mM PMSF, and solubilized for 1 h at 4°C with 1% (w/v) of either n-Dodecyl-β-maltopyranoside (DDM) or lauryl maltose neopentyl glycol (LMNG) (Anatrace), supplemented with 0.2% (w/v) cholesteryl hemisuccinate (CHS, Sigma-Aldrich). After incubation, samples were ultracentrifuged (1 h, 145,000 × *g*, 4°C), and 400 μL of the supernatant was loaded onto a Superose 6 Increase 10/300 GL column (GE Healthcare) equilibrated in 20 mM Tris-HCl pH 7.8, 150 mM NaCl, 10% (v/v) glycerol, and 0.02% (w/v) DDM. Fluorescence-detection size-exclusion chromatography (FSEC) experiments were performed in an ÄKTA^TM^ purifier chromatography system (GE healthcare) coupled to a Jasco FP-4025 fluorescence detector, with GFP fluorescence monitored at 470 nm (excitation) and 510 nm (emission). Chromatograms were recorded and analyzed using UNICORN software.

### Purification of PQLC2 by affinity chromatography using nanoGFP traps

To purifiy the different versions of GFP-tagged PQLC2, we first generated the GFP-trap agarose beads as previously described (48). Briefly, his-tagged anti-GFP nanobodies (nanoGFP, Addgene plasmid # 49172 (66) was expressed in *E. coli* strain *BL21-DE3*, and purified using TALON® metal affinity chromatography. Purified nanoGFP was concentrated and dialyzed overnight at 4°C against PBS to remove imidazole and stored at −20°C with 20% (v/v) glycerol. 4 mL of NHS-Activated Sepharose 4 Fast Flow beads (GE Healthcare Life Sciences) was washed with Milli-Q® H_2_O, activated with 10–15 column-volumes (CV) of 1 mM HCl, and equilibrated with PBS pH 7.5. Then, 4 mg of purified nanoGFP was added to the beads and left overnight at 4°C under gentle rotation, keeping a nanoGFP to beads volume ratio of approximately 0.5–0.7. After the overnight reaction, the remaining non-reacted sites were blocked by the addition of 10–15 CV of 0.1 M Tris-HCl pH 8.5 followed by a 4 h incubation with this buffer at 4°C. Finally, the beads were subjected to 3 cycles of two sequential washes of 3 CV of 0.1 M Tris-HCl pH 8.5, 0.5 M NaCl, and 3 CV of 0.1 M Acetate buffer pH 5.0, 0.5 M NaCl. Finally, the GFP-Trap beads were equilibrated in PBS pH 8.0, and stored at 4°C until use. 20 mL of P3 membranes expressing the different versions of PQLC2-GFP were adjusted to a final protein concentration of 5 mg mL^-1^, and solubilized for 1 h at 4°C in 1% (w/v) DDM and 0.2% (w/v) CHS in buffer containing 20 mM Tris-HCl pH 7.5, 150 mM NaCl, 10 % (v/v) glycerol and 5 mM MgCl₂, and supplemented with 1X PIC, and 1 mM PMSF. Insoluble material was removed by ultracentrifugation (1 h, 145,000 × *g*, 4°C), and the supernatant was incubated for 2 h at 4°C with 4 mL of the nanoGFP trap beads, pre-equilibrated with the column buffer (20 mM Tris-HCl pH 7.8, 150 mM NaCl, 10 % (v/v) glycerol, 0.02 % (w/v) DDM). After incubation, the flowthrough was collected, and the beads were washed with 20 CV of column buffer, followed by 3 CV of elution buffer (20 mM Tris-HCl pH 7.8, 150 mM NaCl, 10 % (v/v) glycerol, 0.02 % (w/v) DDM, 1 mM EDTA, 1 mM DTT). Elution of PQLC2 was performed by incubation at 4°C with 2 CV of elution buffer supplemented with 40 units (40 μg) of Hrv-3C protease (Acro Biosystems). For the experiments of NE-EM, the protein wa eluted after 1 h incubation with the Hrv-3C protease.The eluted fraction was recovered, concentrated using 100 kDa Vivaspin® 20 concentrator (Avantor), and analyzed by Coomassie Blue staining SDS-PAGE. After “in-gel” digestion with chymotrypsin of the SDS-PAGE band atribuited to purified PQLC2, the peptides were analyzed by LC-MSMS triple-TOF at the Platform of Mass Spectrometry (SICaPS) at the I2BC (Gif-sur-Yvette, France).

### Expression and functional assessment of PQLC2 in *Xenopus laevis* oocytes

For oocyte expression, the csPQLC2 cDNA bearing the L290A/L291A mutation was excised from the pYeDP60-csPQLC2-LL/AA-3C-GFP plasmid and subcloned into the pOX(+)-rPQLC2-LL/AA-EGFP plasmid (Jézégou et al., 2012) at the *NcoI* and *BamHI* sites. Capped mRNAs were synthesized *in vitro* from the pOX(+)-rPQLC2-LL/AA-EGFP and pOX(+)-csPQLC2-LL/AA-EGFP plasmids linearized at the *SwaI* site using the mMessage-mMachine SP6 kit (Invitrogen) and stored at −80°C.

Oocytes were prepared from *Xenopus laevis* frogs housed in the local animal facility in compliance with the European Animal Welfare regulations (ethical agreement APAFiS #45643-2023081915049038 v8). Ovarian lobes were extracted from female frogs under anesthesia, and oocyte clusters were incubated on a shaker in OR2 medium (85 mM NaCl, 1 mM MgCl_2_, 5 mM HEPES-K^+^ pH 7.6) containing 2 mg/mL collagenase type II (GIBCO) for 1 h at 25°C. Defolliculated oocytes were sorted under a stereomicroscope and kept at 19 °C in Barth’s solution (88 mM NaCl; 1 mM KCl; 2.4 mM NaHCO_3_; 0.82 mM MgSO_4_; 0.33 mM Ca(NO_3_)_2_; 0.41 mM CaCl_2_; 10 mM HEPES-Na^+^ pH 7.4), supplemented with 50 μg mL^-1^ of gentamycin.

Defolliculated oocytes were injected with 50 ng mRNA (1 µg µL^-1^) encoding rPQLC2-LL/AA-EGFP or csPQLC2-LL/AA-EGFP. Non-injected oocytes were used as negative controls. Two days after cRNA injection, PQLC2 expression at the plasma membrane was monitored under an Eclipse TE-2000 epifluorescence microscope (Nikon) with excitation/emission at 480/535 nm under a 4× objective focused at the equatorial plane. To assay PQLC2 transport activity, oocytes were quickly washed twice with 1 mL ND100 medium (100 mM NaCl, 2 mM KCl, 1 mM MgCl_2_, and 1.8 mM CaCl_2_) buffered at pH 7.5 with 10 mM HEPES-Na^+^. They were then individually incubated for 20 min in 300 µL ND100 medium buffered at pH 5.0 with 10 mM 2-(*N*-morpholino)ethanesulfonic acid (MES)-Na^+^ and supplemented with 0.1 µCi *L*-[2,3,4-^3^H]arginine monohydrochloride (Perkin Elmer; specific activity: 40 to 50 Ci mmol^-1^) and 100 µM unlabelled *L*-Arg. The transport reaction was stopped by three quick washes with 2 mL ice-cold ND100 pH 7.5 medium. Each oocyte was then lysed with 200 µL SDS 10% and mixed with 3 mL EMULSIFIER-SAFE liquid scintillation cocktail (Perkin Elmer). Intracellular radioactivity was counted with a Tri-Carb 2100 TR liquid scintillation analyzer (Packard).

### Negative staining electron microscopy (NS-EM)

Purified csPQLC2 in DDM was concentrated and subjected to size-exclusion chromatography using a Superose 6 Increase 10/300 GL column (GE Healthcare) equilibrated with 20 mM Tris-HCl pH 7.5, 150 mM NaCl, 10% (w/v) glycerol, and 0.015% (w/v) LMNG. Eluted fractions containing csPQLC2 were collected and used directly for NS-EM without further concentration. A 3 μL aliquot of purified protein (∼0.1 mg mL^-1^) was applied to glow-discharged, carbon-coated copper grids and stained twice with 8 μL of 2% uranyl formate. Micrographs were acquired using Talos F200i transmission electron microscope (Thermofisher) operating at 200 kV equipped with a Falcon3 direct electron detector (Thermofisher), at 73000x magnification (pixel size: 1.43 Å). Imaging was performed at the cryo-electron microscopy facility of the I2BC (Gif-sur-Yvette, France) and at the Basque Resource for electron microscopy (BREM) in the Instituto Biofisika (Bilbao, Spain).

### Site-directed spin-labeling (SDSL) of the WDR41-7CD loop

The synthetic single cysteine peptide variants (7CD-A6Px, 7CD-P8Px, 7CD-G20Px, 7CD-V22Px, and 7CDmut-P8Px) of the WDR41-7CD loop (GeneCust) were site-specifically spin-labeled as follows. Peptides (20 μM) were incubated in 25 mM MES buffer pH 7.5 with 10 mM of Tris(2-carboxyethyl)phosphine (TCEP, Alfa Aesar) for 10 min at room temperature to reduce cysteine thiol groups. Labeling was performed by incubating the reduced peptides with 1 mM 3-Maleimido-PROXYL (Sigma-Aldrich) for 1 h at room temperature. To remove TCEP and unreacted 3-Maleimido-PROXYL, the peptide we immovilized in a cation exchange column (Pierce, Thermo Fisher) washed and eluted with 500 mM sodium chloride. To remove the excess of salt, eluted peptides were dyalized in 20 mM Tris-HCl pH 7.5, 150 mM NaCl, 10 % (v/v) glycerol and stored at at −80°C until use.

### EPR spectroscopy measurements and data analysis

EPR spectra were acquired using a Bruker ELEXSYS E500 X-band spectrometer under non-saturating conditions with a modulation amplitude of 1 G. Room-temperature EPR measurements were collected at 294 K using samples loaded into capillary tubes with a final volume of 16 μL. The peptide concentration was maintained at 30-50 μM and csPQLC2 was added at a molar ratio of 1:0.9 (csPQLC2/labeled peptide). For competition assays with the unlabeled peptide, a molar ratio of 1:0.9:5 (csPQLC2/labeled peptide/unlabeled peptide) was used. 50 K EPR spectra were collected using similar peptide and protein concentrations in a final volume of 70 μL. In all cases, the final buffer consisted of 20 mM Tris-HCl, pH 7.5, 150 mM NaCl, 10% (v/v) glycerol, and 0.02% (w/v) DDM.

Room-temperature spectra were simulated using EasySpin (67). Initial values of *A_z_* were set to those obtained from direct measurements of the 50 K spectra for the free and bound peptides. Initial *g* values were chosen according to standard recommendations (68). For multicomponent fitting, the initial parameters of the fast component were set to the values obtained from fitting the spectra of the peptides in the absence of csPQLC2. To obtain a reliable description of *τ*_c_ of the slow component, this component was first fitted independently by masking the spectral region between 3358 and 3410 G. The resulting *τ*_c_ values were then used as initial parameters for the multicomponent fit. Initial values for the intermediate component were set to values intermediate between those of the fast and slow components. The different parameters were allowed to vary as follows: *g_xys_*±0.0002, *A_z_*±4, *A_xy_*±2 and log(*τ*_c_)±0.02 except fot the intermedia for which log(*τ*_c_)±0.2 was used. In all cases a double gaussian/lorentzian line broadening with values of 0.05±0.05 was used.

### Molecular Dynamics Simulations

The initial structure of the csPQLC2/7CD loop complex was generated using the AlphaFold server (v3)(69). When one molecule of csPQLC2 and one of 7CD loop were used as input, AlphaFold positioned the peptide in the putative binding pocket on the cytosolic side of the protein. However, when three molecules of each were used, the peptide was placed on the luminal side. For this reason, monomeric structures of the complex were generated and subsequently aligned into a trimer using the trimeric structure produced by AlphaFold as a template. In this way, a trimeric csPQLC2/7CD loop complex was generated in which the 7CD loop is docked into the cytosolic side of a cytosolic-facing conformation of csPQLC2. This protein complex was then embedded in a DDM micelle composed of 200 molecules using the CHARMM-GUI input generator (https://charmm-gui.org/) (70–73). All simulations were performed using GROMACS version 2024.0 (74,75) with the CHARMM36m force field. The system was placed in a rhombic dodecahedral periodic box extending at least 1.0 nm beyond the longest axis of the complex and solvated with explicit water. The system charge was neutralized and the ionic strength adjusted to 150 mM NaCl. Simulations were carried out in the NPT ensemble at 300 K and 1 atm. Two independent MD simulations of 500 ns each were performed, with atomic coordinates saved every 20 ps.

## Acknowledgement

We want to thank Alice Verchère and Axelle Fillon for their early work and discussions on this project. We also thank Satya Prakash for helping with AF modeling. We thank the animal facility from BioMedTech Facilities, INSERM US36, CNRS UAR2009 at Université Paris Cité for housing the frogs. This work also benefited from the Proteomic-Gif SICaPS, CryoEM, Imagerie-Gif and the Biophysics core facililies of the Institute for Integrative Biology of the Cell (I2BC), supported by the French Infrastructure for Integrated Structural Biology (FRISBI) [ANR-10-INSB-05-05], the Infrastructure for Biology, Health and Agronomy (IBISA), Ile de France Region, Plan Cancer, CNRS and Paris-Sud University. Basque Resource for Electron Microscopy is supported primarily by the Department of Education and the Innovation Fund of the Basque Government, the Fundación Biofísica Bizkaia, and the Spanish Ministry of Science and Innovation, through the Plan de Recuperación, Transformación y Resiliencia (PRTR) funded by the NextGenerationEU (PRTR-C17.I1) program. This work was supported by the Agence Nationale de la Recherche, grant ANR-22-CE11-0008 to BG, JLVI and LT. IU-B was supported by grant PID2022-143177NB-I00 funded by Spanish Ministry of Science, Innovation and Universities.

