## Supplemental Figures 1-8 for "Probing the interaction between the WDR41 7CD loop of the C9ORF72 complex and purified PQLC2"

[1]Université Paris-Saclay, CEA, CNRS, Institute for Integrative Biology of the Cell (I2BC), Gif-sur-Yvette, France; [2]Faculty of Odontology, Université Paris Cité, Oral Health Research Unit, UMR1333, Montrouge, France;[3]Instituto Biofisika (UPV/EHU, CSIC), University of the Basque Country, 48940, Leioa, Spain. [4]Ikerbasque Foundation for Science, Bilbao, Spain, [5]Saints-Pères Paris Institute for the Neurosciences, Université Paris Cité, Centre National de la Recherche Scientifique, F-75006 Paris, France.

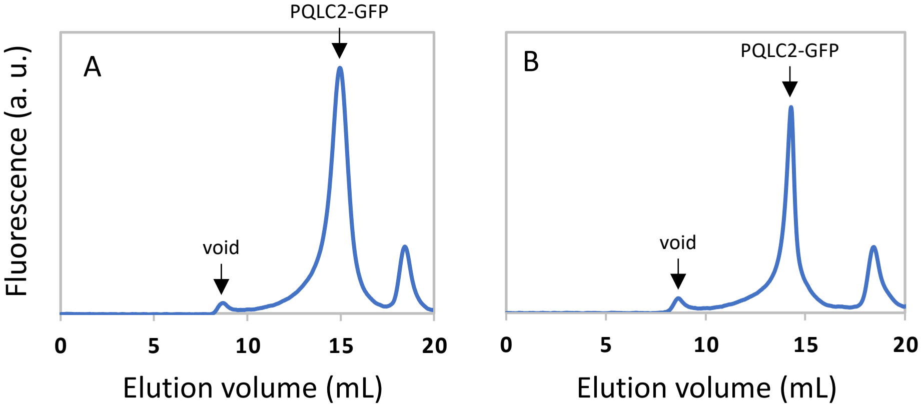


**Supplementary Figure S1. Fluorescence-detection size-exclusion chromatograms of detergent-solubilized LL/AA rPQLC2-GFP.** Membranes containing 5 mg $mL^{-1}$ total protein concentration were solubilized with 2% (w/v) DDM, 0.2% (w/v) CHS (*A*), or 2% (w/v) LMNG, 0.2% (w/v) CHS (*B*), and the supernatant after ultracentrifugation was loaded onto a Superose 6 10/300 GL gel-filtration column equilibrated with 20 mM Tris-HCl pH 7.5, 150 mM NaCl, 10% (v/v) glycerol and 0.02% (w/v) DDM. GFP fluorescence was measured using excitation and emission wavelengths of 470 and 510 nm, respectively. Void volume and PQLC2 are indicated by arrows.

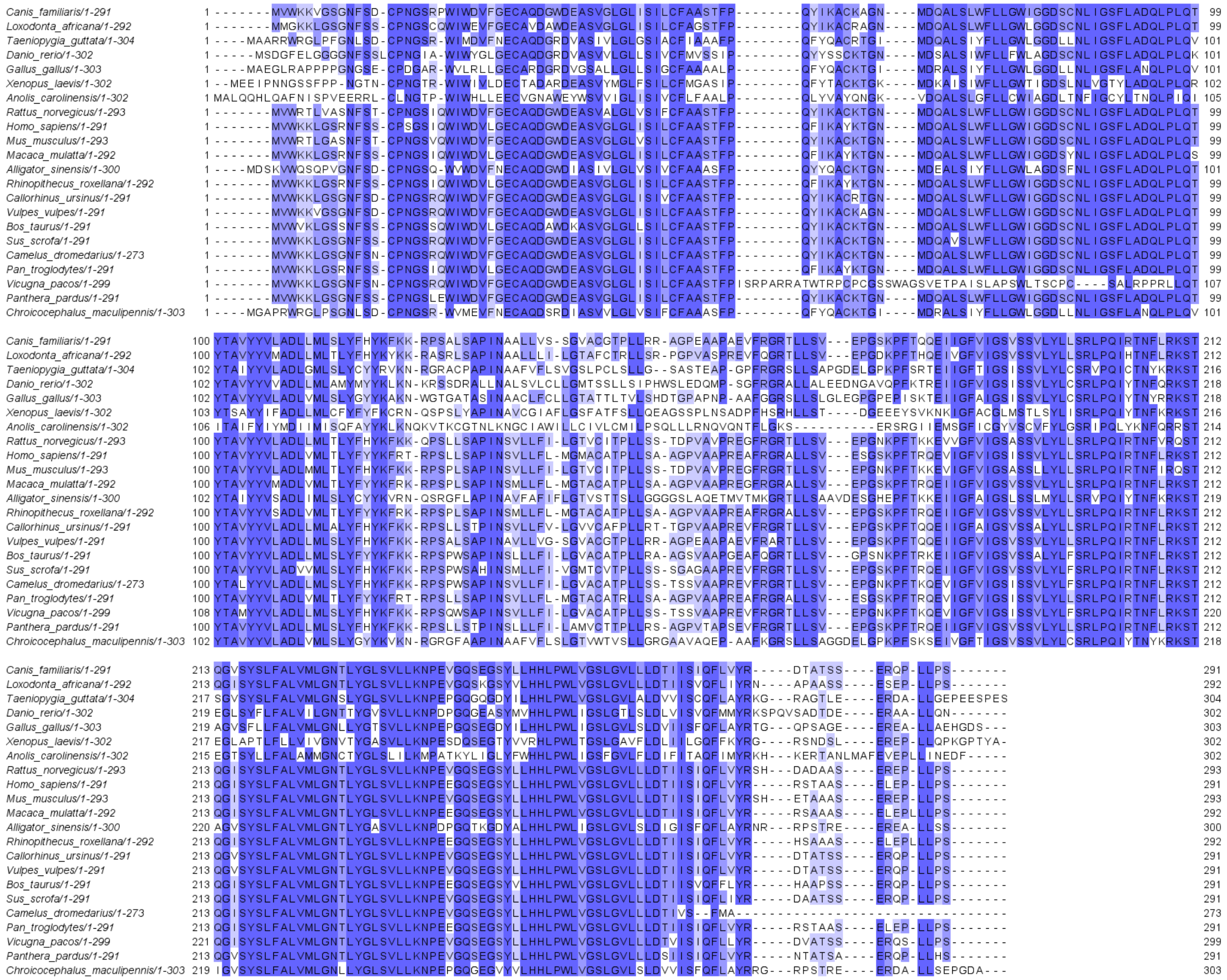

**Supplementary Figure S2**. **Multiple sequence alignment of PQLC2 orthologs**. UniProt accession numbers: *Canis familiaris* (A0A8I3MV82), *Loxodonta africana* (G3T5L2), *Taeniopygia guttata* (A0A674H8J5), *Danio rerio* (A0A8N7UY59), *Gallus gallus* (Q5ZJX0), *Xenopus laevis* (A0A8J1L6L9), *Anolis carolinensis* (R4GCB7), *Rattus norvegicus* (B0BMY1), *Homo sapiens* (Q6ZP29), *Mus musculus* (Q8C4N4), *Macaca mulatta* (A0A1D5QVP4), *Alligator sinensis* (A0A1U7S9B8), *Rhinopithecus roxellana* (A0A2K6R946), *Callorhinus ursinus* (A0A3Q7MW25), *Vulpes vulpes* (A0A3Q7U348), *Bos taurus* (A0A4W2DG10), *Sus scrofa* (A0A4X1W765), *Camelus dromedarius* (A0A5N4DAL8), *Pan troglodytes* (A0A5S6R847), *Vicugna pacos* (A0A6J3B2F5), *Panthera pardus* (A0A6P4WYJ5), *Chroicocephalus maculipennis* (A0A7K5NF94). Residues are colored according to sequence conservation, with darker shades indicating higher conservation. The alignment was performed by Muscle.

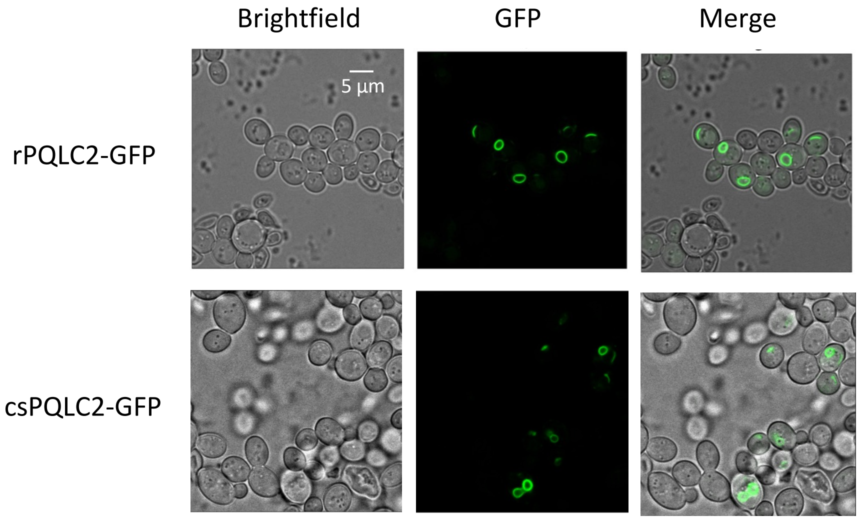


**Supplementary Figure S3**. **csPQLC2-GFP localizes in the vacuole membrane**. Fluorescent confocal microscopy of *S. cerevisiae* cells expressing either rPQLC2-GFP (upper panel) or csPQLC2-GFP (lower panel). The images show that the engineered csPQLC2 expresses at the same membrane compartment than the non-engineered version (rPQLC2).

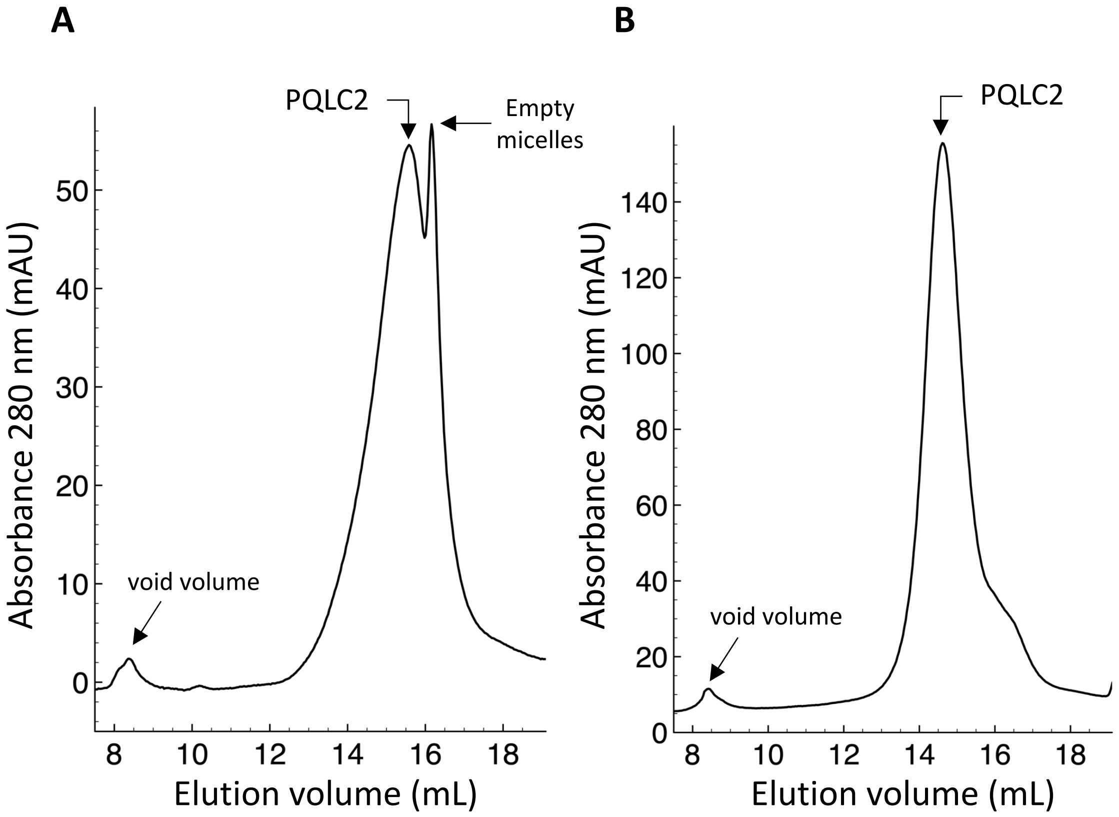


**Supplementary Figure S4. SEC characterization of purified csPQLC2 in detergent solution.** (**A**) SEC profile of purified csPQLC2 in DDM/CHS. DDM/CHS purified protein was concentrated up to 50 µM and loaded onto a Superose 6 increase 10/300 GL column equilibrated with 20 mM Tris-HCl pH 7.8, 150 mM NaCl, 10% (v/v) glycerol, 0.02 % (w/v) DDM, 0.004 % (w/v) CHS. (**B**) SEC profile of purified csPQLC2 in LMNG/CHS. DDM/CHS purified protein was concentrated up to 50 µM and loaded onto a Superose 6 increase 10/300 GL column equilibrated with 20 mM Tris-HCl pH 7.8, 150 mM NaCl, 10% (v/v) glycerol, 0.015 % (w/v) LMNG, 0.003 % (w/v) CHS.

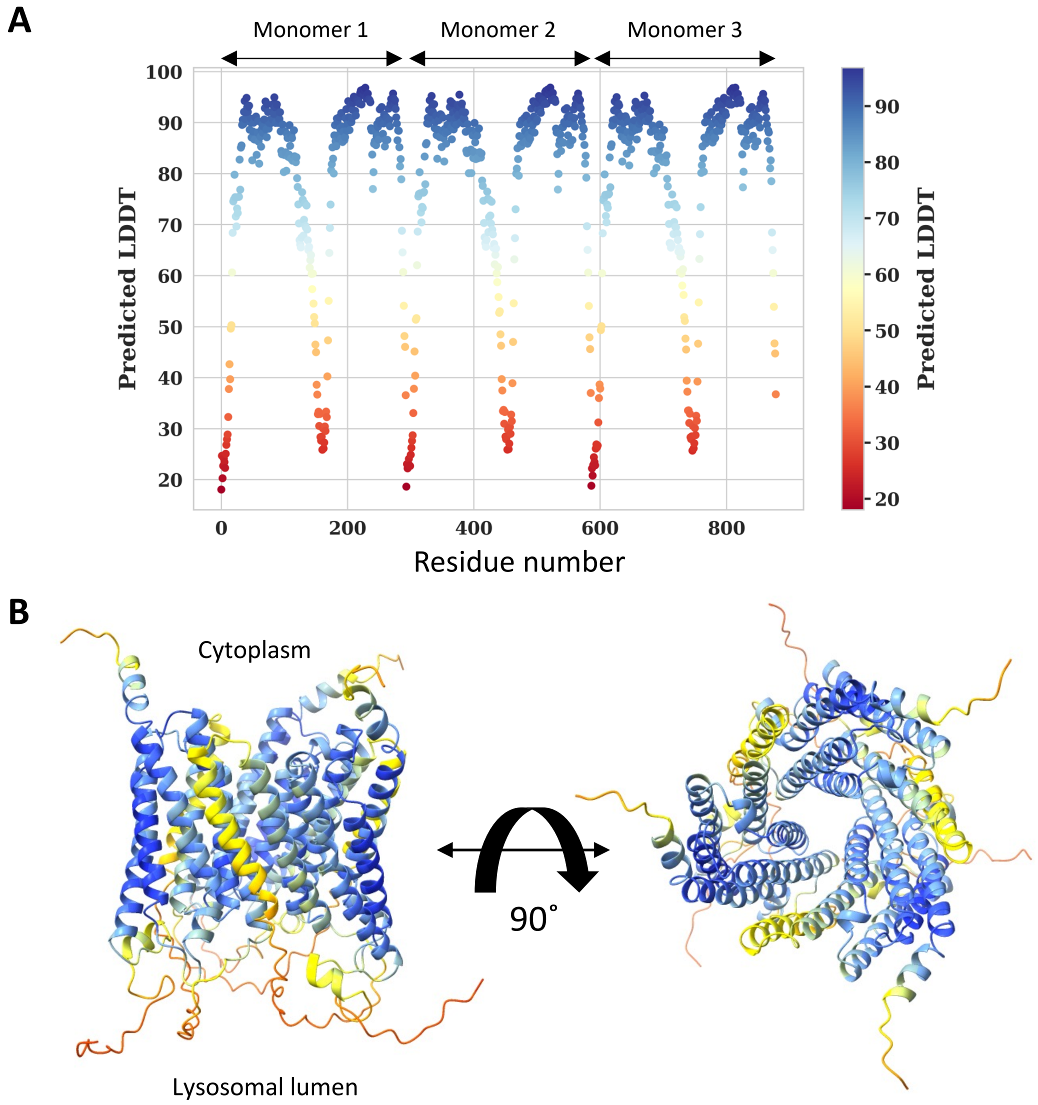


**Supplementary Figure S5. Representation of the pLDDT confident score of the Alphafold trimeric model of rPQLC2**. (A) Predicted LDDT score of each residue of trimeric rPQLC2 in the top-scoring model, colored according to its predicted LDDT value using a gradient from blue (high confidence) to red (low confidence). (B) Cartoon representation of the Alphafold top-scoring trimeric model of rPQLC2 colored as in (A) according the predicted LDDT value at each position. The figure represents the lateral (left) and cytoplasmic-faced (right) orientation.

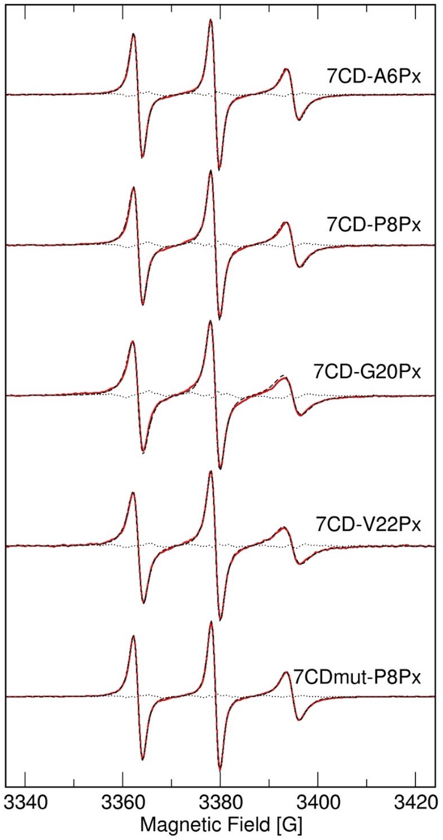


**Supplementary Figure S6. Single-component simulation of free peptides.** Room-temperature, 9 GHz EPR spectra of the various peptides (Figure 6), in buffer containing DDM (red traces) together with the best fits (dashed-black) and the residuals (dotted-black).

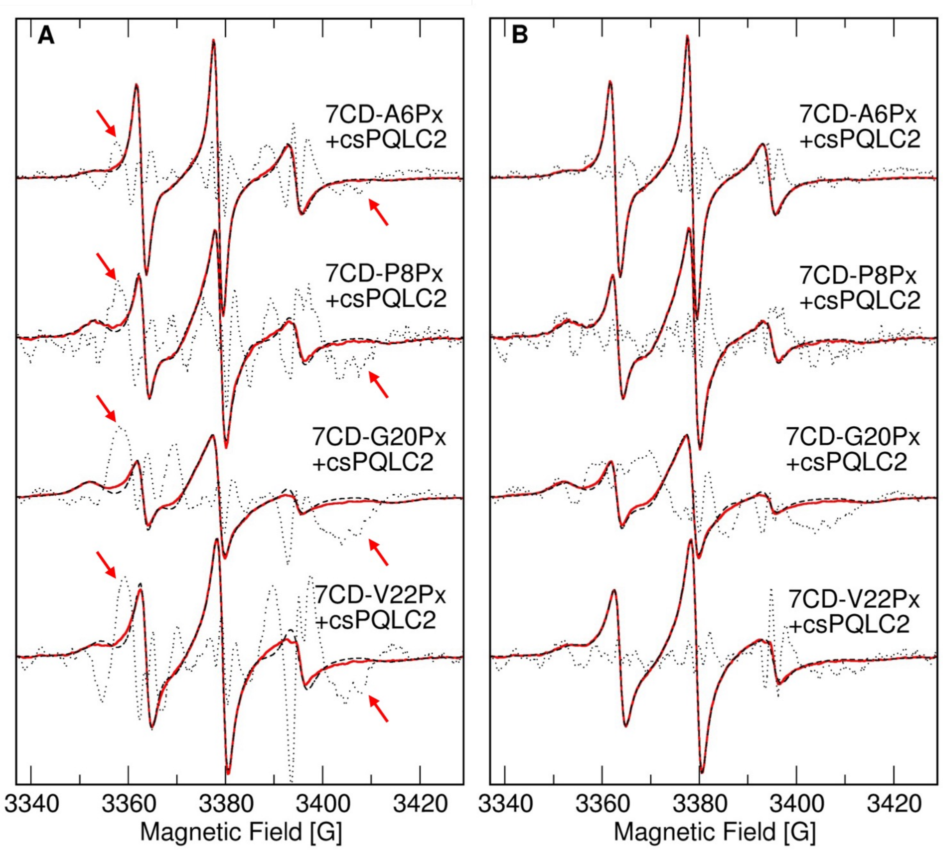


**Supplementary Figure S7. Multiple-component simulation of peptides bound to csPQLC2.** Room-temperature, 9 GHz EPR spectra of the various peptides (Figure 6), in the presence of an equimolar concentration of csPQLC2 (red traces), together with the best fits (dashed black) using a two-component (*A*) or three-component (*B*) model. The residuals (dotted black) have been amplified 10 times to better show the differences. The red arrows indicate peaks that appear in the residuals of the two-component fits, indicative of the presence of a third species with an intermediate rotational correlation time ($\tau_c$).

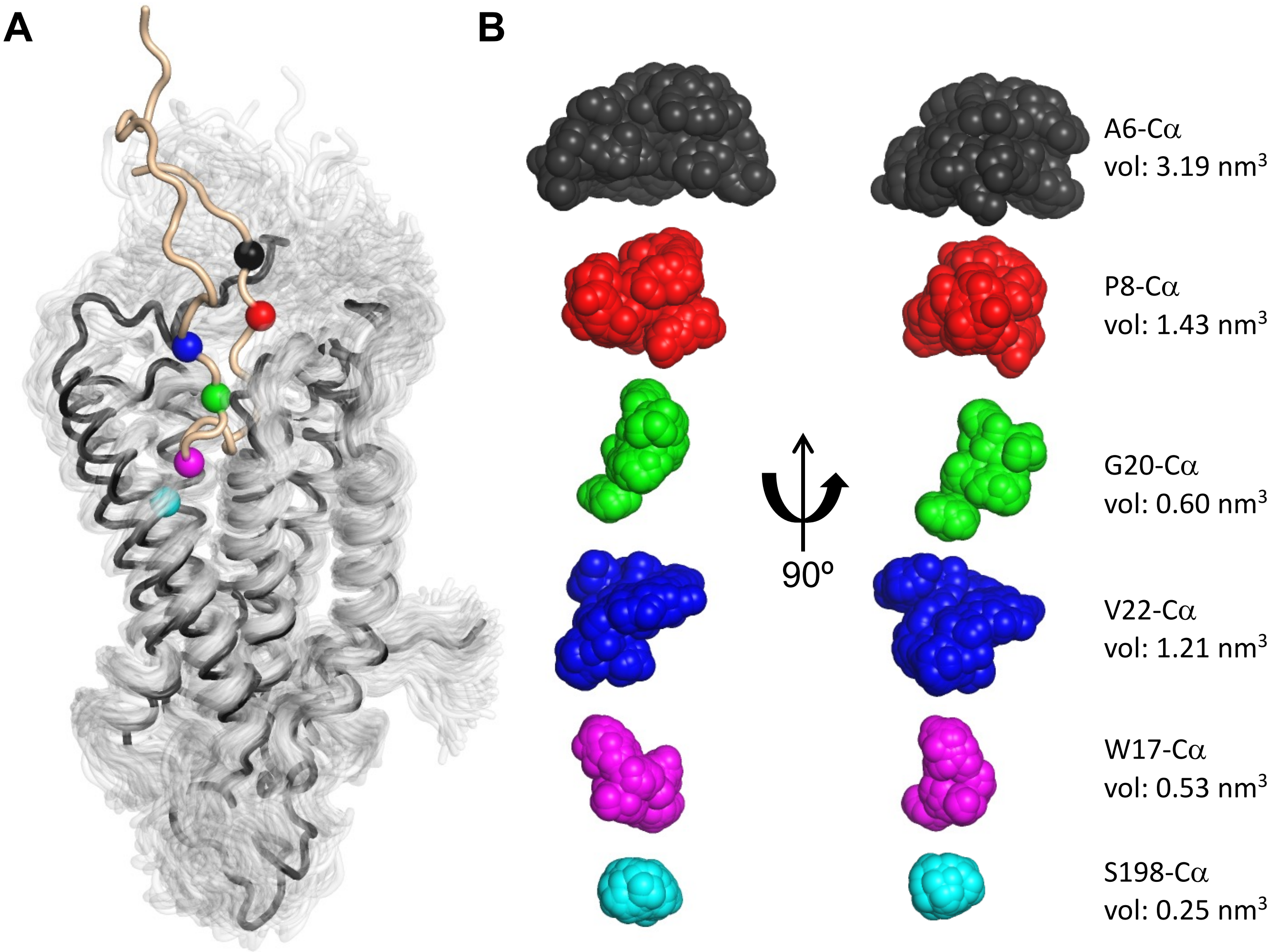


**Supplementary Figure S8. MD simulations of the csPQLC2/7CD loop complex**. (*A*) Initial monomeric structure of csPQLC2 (black) and the 7CD loop (light brown), overlaid with 500 csPQLC2 structures (light grey) sampled at 1 frame/ns along the MD trajectory. Spheres indicate the initial position of the Cα atoms of residues A6 (black), P8 (red), G20 (green), and V22 (blue), used for labeling, as well as residue W17 of the 7CD loop and residue S198 of csPQLC2 (cyan), shown as references. (*B*) Overlay of the Cα positions (same color code as in panel *A*) sampled over the 500 ns simulations at 1 frame/ns for the three monomers from two independent simulations (3000 frames in total). The corresponding explored volumes were calculated as the union of spheres with a radius of 1.5 Å centered on the Cα positions.
